# Recombinant Expression and Purification of *Plasmodium* Heme Detoxification Protein in *E. coli:* Challenges and Discoveries

**DOI:** 10.1101/2025.06.11.659033

**Authors:** Rahul Singh, Ravindra D. Makde

**Affiliations:** Beamline Development and Application Section, Bhabha Atomic Research Centre, Mumbai, India; Homi Bhabha National Institute

**Keywords:** HDP, heme, hemozoin, *Plasmodium*, Soret

## Abstract

Heme, a toxic by-product of *Plasmodium*’s proteolytic digestion of host hemoglobin, is detoxified by the malaria parasite through its conversion into hemozoin (Hz)—the malaria pigment. This detoxification pathway is a key target for many antimalarial drugs, which aim to induce heme-mediated toxicity to the parasite. The Heme Detoxification Protein (HDP) plays a central role in heme-to-Hz transformation; however, its precise mechanism remains unclear, largely due to the absence of successful recombinant expression in a native, soluble form.

In this study, we aimed to express HDP recombinantly in its native soluble state using an *E. coli*-based system. A range of strategies were employed, including expression of orthologs, consensus sequence design, fusion to solubility-enhancing partners, co-expression with molecular chaperones, and extensive construct optimization through N-terminal truncations. Despite extensive efforts, most recombinant HDP constructs were either insoluble or formed soluble aggregates. Notably, only a single construct—with a 44-residue N-terminal truncation and a C-terminal 6His tag (*HDPpf-C10*)—was successfully expressed in a soluble form.

Surprisingly, HDPpf-C10, although retaining domains implicated in heme binding and transformation, exhibited no detectable heme-to-Hz transformation activity. This finding highlights the essential role of the flexible-unstructured N-terminal region in mediating both heme binding and its subsequent conversion to Hz, providing new insights into HDP function and guiding future structural and mechanistic studies.

## 1.3 Introduction

During the asexual multiplication stage of malaria, the *Plasmodium* parasite consumes about 75% of the host’s erythrocyte hemoglobin, providing essential amino acids for its protein synthesis and maintaining osmotic stability [1][2]. This process produces a toxic byproduct, free heme, which can cause cytotoxicity [3]. Unlike other organisms, *Plasmodium* lacks a heme oxygenase pathway for detoxification and instead transforms the heme into harmless hemozoin (Hz) crystals[4],[5]. Antimalarial, drugs interfere with Hz production [6] with chloroquine reported to perturb the nucleation event to induce heme mediated toxicity in malaria parasite [7]. Despite the significance, the specific mechanisms behind Hz crystallization inside parasite food vacuole remain unresolved, and distinct hypotheses have been proposed to explain the rapid production rate of Hz crystals. Deductively, these hypotheses either support spontaneous [8][9] or lipid-assisted Hz crystallization[10], [11][12]. However, a small group of researchers believes Hz crystallization could be a protein-catalyzed process [13],[14][15].

The idea of protein being a potential candidate for mediating catalytic conversion of toxic heme into inert crystalline Hz was put forward in 1992 by A. F. Slater and A. Cerami [13]. Slater et al. identified a novel polymerization/ Hz production activity in the lysate of trophozoites. The Hz formation activity was reported missing in the soluble protein-rich supernatant fraction of the extract and was only present in the plasma membrane and Hz-rich pellet fraction. The Hz formation activity was shown to be inhibited by SDS exposure. The authors concluded that this Hz formation activity might be due to heme or lipid binding protein, explaining the presence of activity only in the pellet fraction of trophozoite lysate.

Four years later (1996), after discovering Hz formation activity in trophozoites lysate, the first actual protein candidate was presented to the world by a group led by Daniel E. Goldberg [14]. The *plasmodia* histidine-rich protein (HRP) shows a high binding affinity to heme and, in in-vitro conditions, can catalyze free heme’s conversion into Hz. Interestingly, Hz production is unanimous among all species of *Plasmodium*. However, HRP is only present in the *falciparum* species of *Plasmodium* and is missing in other species like *vivax*, *malaria*, and *ovauole*. This observation raised concerns about HRP being the actual biological catalyst for Hz production. However, this study acts as a proof of concept that a protein can catalyze the transformation of heme into Hz.

The next breakthrough in this area came in 2009 when a new and highly conserved *Plasmodial* protein known as heme detoxification protein (HDP; UniProt ID: Q8IL04) was identified in *P. falciparum* [15]. HDP was found to be most efficient in heme to Hz transformation among all known mediators of Hz production. The importance of *HDP* gene for *Plasmodium* survival is underscored by the inability to isolate parasite strains with mutations in the HDP gene despite repeated attempts, suggesting that such mutations are likely lethal or severely compromise parasite viability [15] [16]. Because of its crucial role in the physiology of malaria parasites, HDP became a topic of enormous interest among malariologists. In recent years, several food vacuole proteases (falciparum-2, plasmepsin-2, plasmapsin-4, and histo-aspartic proteases) and HDP were co-purified as a complex from parasites [17], suggesting that these proteins interact in vivo and form associations that may facilitate hemoglobin proteolysis and transfer the liberated heme group to HDP for assembly into Hz.

Despite the pivotal role the HDP protein plays in malaria pathogenesis and its potential significance in antimalarial drug development, the precise workings of the HDP gene have remained enigmatic. This knowledge gap can be attributed to previous studies focusing on HDP in a detergent-solubilized state extracted from inclusion bodies rather than investigating the native protein. To bridge this gap, our study aimed to achieve the recombinant expression of *Plasmodium* HDP in its native soluble form through an *E. coli*-based bacterial expression system. Employing various strategies, we successfully obtained a truncated version of HDP expressed in a native soluble state. Our investigation into this soluble protein unveiled the pivotal role of the first 44 flexible-unstructured residues of HDP in governing both heme binding and transformation activity.

## 1.4 Material and method

### 1.4.1 Material

Flat bottom, 96-well plates were purchased from Corning (3635). Dimethyl sulfoxide (DMSO), hemin (Sigma-Aldrich, ≥97%, 51280-1G), sodium acetate and Tris base were obtained from Sigma-Aldrich. Glacial acetic acid and acetone were procured from EMPURA.

### 1.4.2 Cloning

Putative homologs of the Heme detoxification protein (*hdp)* gene were identified through BLAST analysis by using the HDP sequence from *Plasmodium falciparum* (HDPpf; UniProt ID: Q8IL04) as a search template. Homologous coding DNA sequences (CDS) were identified for *hdp* genes from *Plasmodium vivax* (UniProt ID: A0A1G4HHQ7), *Plasmodium knowlesi* (UniProt ID: A0A384L7P6), *Theileria equi* (UniProt ID: L0B1I2), and *Babesia bigemina* (UniProt ID: A0A061DBU8). These genes were codon-optimized and commercially synthesized. The optimized CDS of all four *hdp* genes were cloned between the BamHI and HindIII restriction sites of pST50Tr, a T7-promoter-based expression plasmid, to form an in-frame translational fusion with streptavidin-His6-TEVtag (STRHISTEV, 3.8 kDa). The accuracy of all constructs was verified by DNA sequencing.

### 1.4.3 Preparation of consensus design cassette of HDP

Consensus design approach, has been shown earlier to improve recombinant protein solubility, though increasing thermostability and folding robustness [18]. To prepare the consensus design of *HDP* gene (*HDPcc*), the amino-acid conservation of 17 homologs of HDP from different species of *Plasmodium* (*P. falciparum, P. berghei, P. chabaudi, P. coatneyi Hackeri, P. yoelii, P. vivax, P. vinckei, P. relictum, P. reichenowi, P. ovale curtisi, P. malariae, P. knowlesi, P. inui, P. gallinaceum, P. gaboni, P. fragile, P. cynomolgi*) were analyzed by multiple sequence alignment **(*Figure S1*)**. The amino acid conservation at each position was manually analyzed to prepare a consensus/ average conserved sequence of HDP. The gene coding this consciences sequence was commercially synthesized. HDPcc gene was cloned between BamHI and HindIII restriction site of pST50Tr vector to form an in-frame straptavedin-6His-TEV-HDPcc fusion protein.

### 1.4.4 Bioinformatic analysis

Disordered region in HDPpf protein were identified by using DISOPRED3 [19] and Ab initio 3D model of HDPpf was prepared by using AlphaFold 3 [20]. The structure space homology was determined via DALI [21]

### 1.4.5 Construct optimization

Four constructs of HDPpf gene namely HDPpf-N1, HDPpf-N2, HDPpf-N3 and HDPpf-N4 with deletion of 11, 21, 27, 37 N-termini residue respectively and 3 C-terminal residues were cloned between BamHI and HindIII restriction site of pST50Tr, a T7-promoter based expression plasmid, to form in-frame translational fusion with streptavidin-His6-TEVtag (***Figure S2*A**; ***Table 1***). Similarly, amplicons with various truncations at N-terminus of HDPpf (HDPpf-C1 to HDPpf-C10) were cloned between NcoI & XhoI restriction site in pET28a, an *E.coli* protein expression vector with C terminal 6-His tag (***Figure S2*B**; ***Table 1***). HDPpf-C10 (Δ44N) equivalent truncated amplicons of *hdp* gene, from *Babesia bigemina* (Δ39N; HDPbbC1), *Theileria equi* (Δ39N; HDPtqC1), *Plasmodium vivax* (Δ44N; HDPpvC1) and *Plasmodium knowlesi* (Δ58N; HDPpkC1) was cloned between NcoI & XhoI restriction site in pET28a, with C-terminal 6-His tag. The accuracy of all constructs was verified by DNA sequencing.

**Table 1.** HDP constructs, properties, and recombinant expression.

| Sr. No. | Construct | Organism | Vector | Truncation |  | N-terminal Fusion protein | C-terminal Fusion protein | Molecular weight (kDa) | pI | Expression Temperature (°C) |  |  | Co expression - GroEI | Co expression - DnaK | Remark |
| --- | --- | --- | --- | --- | --- | --- | --- | --- | --- | --- | --- | --- | --- | --- | --- |
|  |  |  |  | N-terminal truncation | C-terminal truncation |  |  |  |  | 37 | 28 | 37 / 16 |  |  |  |
| 1. | HDPpf | *P. falciparum* | pST50Tr | NA | NA | N-STR-6His-TEV (3.796 kDa) | NA | 28.13 | 9.17 | P | P | P | P | S* | Expressed in Rosetta(DE3)pLysS |
| 2. | HDPpv | *P. vivax* | pST50Tr | NA | NA |  | NA | 27.74 | 9.75 | P | P | P* | Not determined (N.D.) |  |  |
| 3. | HDPcc | Consensus Sequence | pST50Tr | NA | NA |  | NA | 27.93 | 9.08 | P | P | P | P | P |  |
| 4. | HDPpk | *P. knowlesi* | pST50Tr | NA | NA |  | NA | 25.57 | 9.61 | P | P | P* | N.D. |  |  |
| 5. | HDPbb | *Babesia bigemina* | pST50Tr | NA | NA |  | NA | 26.61 | 9.49 | P | P | P* | N.D. |  |  |
| 6. | HDPtq | *Theileria equi* | pST50Tr | NA | NA |  | NA | 26.64 | 9.75 | P | P | P* | N.D. |  |  |
| 7. | HDPpf-N1 | *P. falciparum* | pST50Tr | 11 | 3 |  | NA | 26.34 | 8.97 | P | P | P | P | P |  |
| 8. | HDPpf-N2 | *P. falciparum* | pST50Tr | 21 | 3 |  | NA | 25.32 | 8.33 | P | P | P | P | P |  |
| 9. | HDPpf-N3 | *P. falciparum* | pST50Tr | 27 | 3 |  | NA | 24.47 | 6.97 | P | P | P | P | P |  |
| 10. | HDPpf-N4 | *P. falciparum* | pST50Tr | 37 | 3 |  | NA | 23.31 | 6.97 | S* | S* | S* | N.D. |  |  |
| 11. | HDPpf-C1 | *P. falciparum* | pET28a | NA | NA | NA | 6His | 25.5 | 9.19 | P* | P* | P* | p | p | Shown expression only in C41 and C43 (DE3) |
| 12. | HDPpf-C2 | *P. falciparum* | pET28a | 8 | NA | NA | 6His | 24.5 | 9 | P | P | P | p | p |  |
| 13. | HDPpf-C3 | *P. falciparum* | pET28a | 18 | NA | NA | 6His | 23.2 | 8.35 | P | P | P | S* | S* |  |
| 14. | HDPpf-C4 | *P. falciparum* | pET28a | 22 | NA | NA | 6His | 22.8 | 7.82 | P* | P* | P* | p | S* | Shown expression only in C41 and C43 (DE3) |
| 15. | HDPpf-C5 | *P. falciparum* | pET28a | 28 | NA | NA | 6His | 22.2 | 6.95 | P | P | P | S* | S* |  |
| 16. | HDPpf-C6 | *P. falciparum* | pET28a | 25 | NA | NA | 6His | 22.3 | 7.27 | P* | P* | P* | S* | S* | Shown expression only in C41 and C43 (DE3) |
| 17. | HDPpf-C7 | *P. falciparum* | pET28a | 31 | NA | NA | 6His | 21.8 | 7.27 | P | P | P | p | S* |  |
| 18. | HDPpf-C8 | *P. falciparum* | pET28a | 34 | NA | NA | 6His | 21.5 | 7.82 | P | P | P | p | S* |  |
| 19. | HDPpf-C9 | *P. falciparum* | pET28a | 37 | NA | NA | 6His | 21.1 | 6.95 | P | P | P | S* | S* |  |
| 20. | HDPpf-C10 | *P. falciparum* | pET28a | 44 | NA | NA | 6His | 20.3 | 7.28 | P | <u>S</u> | <u>S</u> | N.D. |  |  |
| 21. | HDPpv-C1 | *P. vivax* | pET28a | 44 | NA | NA | 6His | 19.82 | 7.38 | N.D. |  | S* | N.D. |  |  |
| 22. | HDPpk-C1 | *P. knowlesi* | pET28a | 58 | NA | NA | 6His | 19.4 | 8.51 | N.D. |  | <u>S</u> | N.D. |  |  |
| 23. | HDPbb-C1 | Babesia bigemina | pET28a | 39 | NA | NA | 6His | 19.28 | 6.97 | Not tried |  | <u>S</u> | N.D. |  |  |
| 24. | HDPTqC1 | Theileria equi | pET28a | 39 | NA | NA | 6His | 19.38 | 8.53 | Not tried |  | <u>S</u> | N.D. |  |  |
| 25. | HDPpf-GFP | P. falciparum | pET33b | NA | NA | NA | TEV-GFP-6His (28.4kDa) | 53.3 | 7.39 | P | P | P | P | P | Expressed in C41 (DE3) |
| 26. | TRX-HDPpf | P. falciparum | pST50Tr | NA | NA | 6His-TRX-TEV (15.2kDa) | NA | 39.38 | 8.44 | P | P | P | P | P |  |
| 27. | TRX-HDPcc | Consensus sequence | pST50Tr | NA | NA |  | NA | 39.18 | 8.13 | P | P | P | P | P |  |
| 28. | MBP-HDPpf | P. falciparum | pST50Tr | NA | NA | 6His-MBP (Small-linker) (35.7 kDa) | NA | 66.8 | 7.18 | P | P | P | S | S |  |
| 29. | pCold-HDPpf | P. falciparum | pCold<sup>TM</sup>TF | NA | NA | pCold-TF (52.18kDa) | NA | 80.27 | 6.39 | Not Applicable |  | <u>S*</u> | N.D. |  | Expression @ 16°C |
| 30. | pCold-HDPcc | Consensus sequence | pCold<sup>TM</sup>TF | NA | NA |  | NA | 80.07 | 5.99 |  |  | <u>S*</u> | N.D. |  |  |

**P:** Expression in pellet fraction;
**P\*:** Expressed protein is primarily in pellet fraction but small amount is in supernatant fraction which is not responsive for Ni-IDA chromatography and lacks heme-to- Hz transformation activity;
**S\*:** Expressed protein is significantly present in soluble supernatant fraction but non responsive for Ni-IDA chromatography, and lacks heme-to-Hz transformation activity;
**NE:** Not expressed;
**S:** Expression in soluble phase and protein responsive to Ni-IDA but but lacking heme-to-Hz transformation activity.
Not
determined

### 1.4.6 Cloning of Fusion protein constructs

a. *GFP fusion Protein Construct:* An amplicon of *GFP* with N-terminal TEV protease site was first cloned at Kpn1 and Sac1 restriction site of pET33b(+) vector. Full length amplicon of HDPpf was then later cloned between NdeI and kpn1 restriction site of this modified vector to yield *HDP-TEV-GFP-6His* fusion protein construct (***Figure S*3A**).
b. *Thyroidin fusion Protein Construct:* The genes encoding thioredoxin (*TRX*; UniProt ID: P0AA25; *E. coli*) was amplified, introducing NHE1 restriction sites at both the 5’ and 3’ ends of the amplicons. Site-directed mutagenesis was employed to create an NHE1 restriction site between the STR-6HIS sequence and TEV-MCS in the pST50Tr vector. Subsequently, the amplicons were cloned into this modified vector. The alignment of the cloned *TRX* gene was confirmed through PCR, protein expression studies, and sequencing. Following this, HDPpf was cloned between the BamHI and HindIII restriction sites of the modified pST50Tr vector, resulting in the formation of the respective fusion protein constructs (***Figure S3*B**).
c. *Maltose binding protein (MBP) fusion protein construct*: The MBP tag was amplified via PCR, from pHLIC-MBP vector with 5’ NdeI and 3’ BamHI restriction site. This amplicon also has nucleotide coding for TEV protease cleavage site between C-termini of *MBP* coding region and BamHI restriction site at site at 3’ end. This amplicon was later cloned NdeI and BamHI site of pST50Tr vector with HDPpf **(*Figure S*3C)**.
d. *pCold trigger factor fusion Protein Construct* (***Figure S*3D**): The *HDPpf* and *HDPcc* construct between NdeI, and HindIII restriction enzyme was sub-cloned from pST50Tr vector (Section 1.4.2) to pCold TF DNA Vector (Takara). This sub-cloning resulted in a fusion protein construct with an N-terminal pColdtrigger factor (48kDa) followed by 3C-thrombin-factor Xa precision protease site and Straptividin 6His-TEV-HDP at C-terminus **(*Figure S3* D)**.

### 1.4.7 Recombinant protein expression and purification Expression

All construct were transformed into *E. coli* strains [BL21(DE3)/ BL21(DE3) pLysS/ C41(DE3)/ C43(DE3)]. A single isolated colony was inoculated into 50ml of 2XTY media with ampicillin (50 µg/ml) or kanamycin (30 µg/ml) and/ or chloramphenicol (34 µg/ml). The culture was allowed to grow at 37 °C under shaking condition till OD at 600 nm reached 0.8 and then the temperature was shifted to 18°C/28 °C. After half hour of acclimatization at shifted temperature, the culture was induced to express recombinant protein by adding 0.2 mM Isopropyl β-D-1-thiogalactopyranoside. The culture was then allowed to grow on 37°C for 4 hours, 28 °C for 12 hours or 18°C for 24 hours. Mature culture was harvested by centrifugation at 14,300 g (RCF) for 5 minutes. Cell pellet was re-suspended in “20 mM Tris-Cl pH 8, 400 mM NaCl” buffer. Re-suspended cells were frozen in liquid nitrogen followed by thawing at 30°C. Slurry homogenization was performed on ice using probe sonicator. Homogenized slurry was centrifuged at 42000 g (RCF) at 4°C for 30 min. Supernatant was used for protein purification by immobilized metal affinity chromatography (IMAC) using Ni-IDA resin (GE).

### 1.4.8 Screening of solubilizing chaperones

The required construct was transformed in *E. coli* BL21(DE3) or C41(DE3) or C43(DE3). The transformed cells were further co-transformed with pG-TF2, pKJE7, pG-KJE8, pTF16, pGro7 plasmids (TAKARA Cat. #3340). The 100ml 2xTY media was inoculated with single isolated recombinant colony of each of the 5 class of transformed cells. All 5 cultures were allowed to grow at 37 °C under shaking condition till OD at 600 nm reached 0.6 and then the temperature was shifted to 18 °C and first induction to express chaperone proteins, was given by adding 5 µg/ml of L-arabinose. Culture was allowed to grow at 18 °C till OD at 600 nm reached 0.8 and then induced by adding 0.2 mM isopropyl β-D-1-thiogalactopyranoside. The culture was then allowed to grow for 24 hours on 18 °C. Cells were harvested by centrifugation at 14,300g for 5 minutes. Cell pellet was suspended in 20 mM Tris-Cl buffer at pH 8 supplemented with 400 mM NaCl and1mM MgCl_2_. Suspended cells were flash frozen in liquid nitrogen followed by thawing at 30 °C in presence of lysozyme. Slurry homogenization was performed at 4°C using probe sonicator. Slurry was centrifuged at 4 °C for 10 min. Pellet was discarded and supernatant was used for protein purification by immobilized metal affinity chromatography (IMAC) using Ni-IDA resin (GE Healthcare). 1mM MgCl_2_ is maintained during all steps of down-stream processing, purification and protein concentration.

### 1.4.9 Expression of HDP as a fusion protein with N-terminal pCold trigger protein tag

The required construct (vector pCold^TM^TF) was transformed in *E. coli* BL21 (DE3)pLySS. The transformed cells were allowed to grow at 37 °C under shaking condition till OD at 600 nm reached 0.7 and then the temperature was shifted abruptly to 14 °C by incubating the culture flask to 14 °C water bath for half an hour. After this cold shock, the culture is induced by adding 0.2 mM isopropyl β-D-1-thiogalactopyranoside. The induced culture was then allowed to grow for 36 hours on 14 °C to maturation. The mature culture is harvested and processed in the same way as mentioned above.

### 1.4.10 Inclusion body purification

Isolated inclusion bodies, obtained after centrifugation, were resuspended by sonication in a solution composed of 1M NaCl and a 50mM NaHCO_3_ buffer at pH 10. The homogenized inclusion bodies were subsequently subjected to centrifugation at 42,000g (RCF) for 30 minutes. The resulting supernatant was discarded, and the pellet underwent 3 to 4 washes using a buffer containing 50mM NaHCO_3_ at pH 10. In each washing step, the pellet was homogenized in the solution and then centrifuged at 42,000g (RCF) for 10 minutes. Finally, the purified inclusion bodies were homogenized in a 20% (V/V) glycerol solution and stored at −80°C for further use.

### 1.4.11 Inclusion body solubilyzation via N-Lauryl Sarcosine

Purified inclusion bodies were solubilized in a solubilization buffer containing 50 mM CAPS (pH 11.0), 1.5% N-lauryl sarcosine and 0.3 M NaCl, for 30 minutes. The solubilized protein was separated by centrifugation and purified via metal immobilization chromatography. The solubilized protein was allowed to bind with Ni-IDA resin pre-equilibrated with a solubilization buffer. To remove non-specifically bound protein, the resin was washed with a buffer containing 50 mM CAPS pH 11.0, 1M NaCl, 0.3% N-lauryl sarcosine and 50 mM imidazole. The specifically bound protein was eluted via imidazole gradient (20 to 500 mM) in a buffer containing 50 mM CAPS pH 11.0, 0.3% (V/V) N-lauryl sarcosine and 0.3 M NaCl. Before the enzymatic assay, the impurities like imidazole and N-lauryl sarcosine were removed from purified HDP protein via desalting chromatography with column (GE HiTrap Desalting 5ml) equilibrated with a buffer containing 10 mM CAPS pH 11 and 135 mM NaCl.

### 1.4.12 Small-angle X-ray scattering (SAXS) data processing and modeling

The small-angle X-ray scattering (SAXS) data for the DnaK-HDPpf complex were collected using an Anton Paar SAXSpace instrument equipped with a Copper Kα laboratory source and the data was collected with default instrument parameters.

Data analysis and modeling on data sets were predominantly performed using software tools available in the ATSAS suite[22]. The radius of gyration (Rg) and absolute intensity I(0) at q = 0 were estimated from the pair distance distribution [P(r)] calculated using GNOM from the ATSAS suite. The molecular weight of the protein was estimated using a concentration-independent porod invariant estimator based on Bayesian inferences and machine learning on shape-categorized SAXS data implemented in BioXTAS RAW software[23]. As the SAXS data were on an arbitrary relative scale, fitting of solution scattering from macromolecules with known atomic structure to the experimental scattering curves was carried out using the CRYSOL tool [24] in the ATSAS package. To generate ab-initio Dummy chain models, GASBOR[25], with no constraints for the DnaK-HDPpf complex. The experimental data has been submitted to SASBDB database.

### 1.4.13 Pyridine hemchrome assay

The assay was performed as described earlier [26].

### 1.4.14 Heme Binding Assay

The heme binding properties of the HDP protein were assessed using spectroscopic titration assays. Purified HDP (either HDPpf or HDPpf-C10) with a histidine tag was buffer-exchanged into a solution containing 1 mM imidazole, 150 mM NaCl, and 20 mM Tris–HCl at pH 8. This buffer exchange was accomplished using a HiTrap desalting column (5 mL; Citiva) packed with Sephadex G-25 Superfine resin. A monomeric hemin stock solution was prepared following a previously described method[27]. Initially, hemin chloride was dissolved in 500 μL of 0.1 M NaOH, followed by the addition of 500 μL of 1 M Tris (pH 8.0). The concentration of the stock solution was determined spectrophotometrically, utilizing an extinction coefficient of 58.4 mM^−1^ cm^−1^. This stock solution was diluted in the buffer (1 mM imidazole, 150 mM NaCl, and 20 mM Tris–HCl, pH 8) to reach a final concentration of 80 µM. Successive aliquots of 10 µM hemin, combined with varying concentrations of HDP (ranging from 0 to 18 μM), were incubated at room temperature for 30 minutes. Equal volumes of 180 µL from each reaction were loaded into a 96-well plate, and UV/Vis spectra were recorded between 300 and 700 nm using a BMG Labtech POLARstar Omega Microplate Reader at 25°C. To determine the K_d_ value, the difference in absorbance at 410 nm (410nmΔA), i.e., absorbance at 410 nm with X μM HDP (X = 0 to 16 μM), subtracted from the absorbance at 410 nm with 0 μM HDP, was plotted against HDP concentration. The data were fitted to the Langmuir binding isotherm equation (Equation1) to determine the Kd value, following the formula:

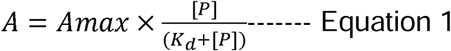

*Where: A is the absorbance at 410 nm; Amax is the maximum change in absorbance (theoretical maximum value); [P] is the concentration of the protein. K_d_ is the dissociation constant to be calculated*.

## 1.5 Results

### 1.5.1 Overview of Expression Strategy

To achieve soluble recombinant expression of the Heme Detoxification Protein (HDP), we employed multiple strategies: homolog expression, fusion tags, N-terminal truncations, consensus sequence design, chaperone co-expression, and optimization of expression conditions. Constructs were assessed for expression level, solubility, responsiveness to affinity purification, and heme-to-Hemozoin (Hz) transformation activity.

Initial expression of full-length HDPpf and its homologs (*HDPpv*, *HDPpk*, *HDPbb*, *HDPtq* and *HDPcc*) at 37°C, 28°C, and 18°C yielded high levels of recombinant protein. However, SDS-PAGE analysis showed that the majority of protein was confined to the insoluble pellet fraction. At 18°C **(Figure S4)**, faint bands corresponding to the expected molecular weight were observed in the supernatant for all constructs except *HDPbb* **(Figure S4E)**, indicating limited soluble expression. These soluble fractions failed to bind Ni-IDA resin, suggesting that the proteins were either misfolded or present as soluble aggregates. In addition, none of the supernatant or bound fractions exhibited detectable heme-to-Hz transformation activity.

### 1.5.2 Fusion Protein Strategies Improve Solubility but Not Activity

To enhance soluble expression of HDPpf, several fusion tags—GFP, MBP, TRX, and pCold Trigger Factor—were tested under low-temperature expression (18°C). The TRX-HDPpf fusion, despite thioredoxin’s known solubilizing properties, showed poor expression and remained largely insoluble across all temperatures (18°C, 28°C, and 37°C). The HDPpf-GFP fusion expressed in C41(DE3) produced visible fluorescence, but most protein localized to the insoluble fraction. Soluble fluorescence was attributed to cleaved GFP rather than intact fusion protein, and Ni-IDA chromatography primarily recovered free GFP **(Figure S5A, B)**. The MBP-HDPpf fusion, when expressed at 18°C, showed significant soluble protein. However, the protein failed to bind Ni-IDA or ion-exchange columns, suggesting it existed as soluble aggregates. Finally, the pCold-HDPpf fusion, expressed via a cold-shock promoter system, yielded soluble protein at 14°C **(Figure 1A)**. However, biophysical studies indicated the presence of aggregates. While thrombin—but not TEV—could cleave the fusion, the released HDP precipitated upon separation, indicating misfolding **(Figure 1B)**. Together, these results show that although fusion tags improved HDPpf solubility, none of the constructs displayed heme-to-Hz transformation activity, likely due to improper folding.

**Figure 1.**
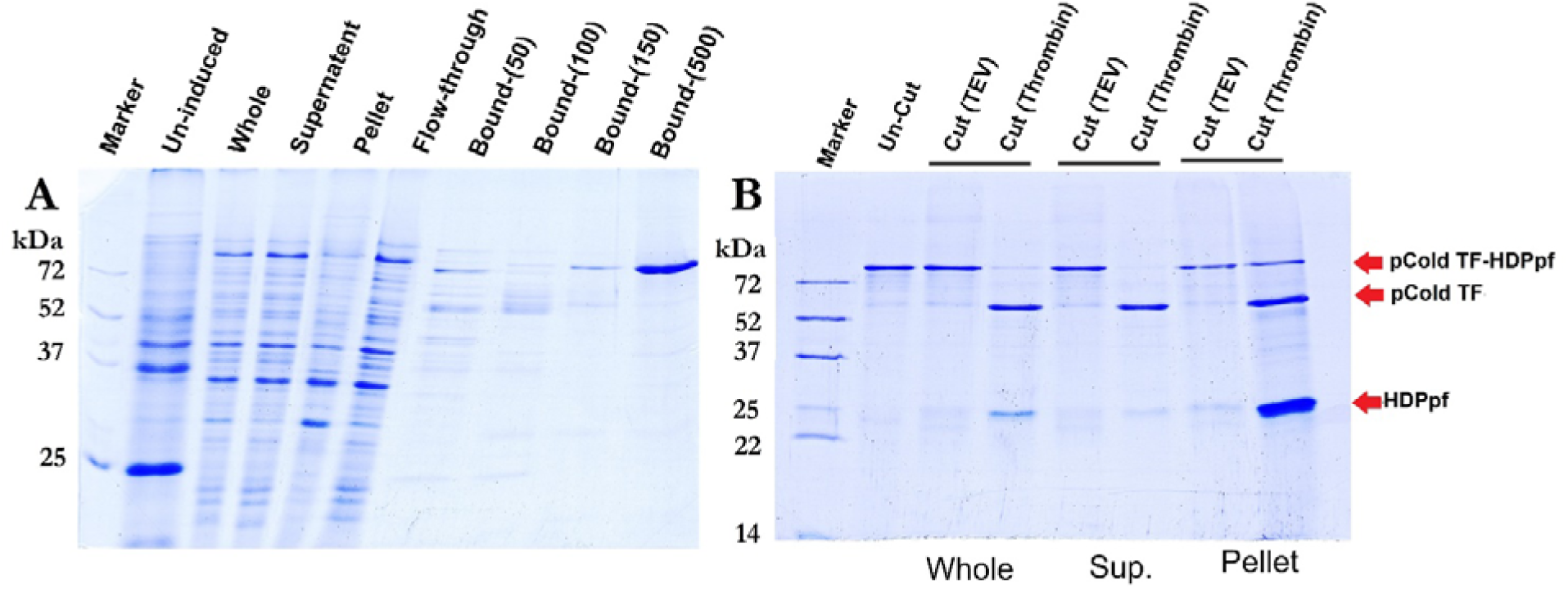
Coomassie Blue-stained SDS-PAGE (15% w/v) gel of (A) various samples collected during the expression of pCold-HDPpf fusion protein construct. **B,** the digestion of soluble recombinant pCold-HDPpf fusion protein with thrombin and TEV precision proteases where whole sample correspond to all of the sample whereas Sup. and pellet are the fractions collected after the centrifugation of whole sample.

### 1.5.3 Co-expression with Chaperones Facilitates Solubility

Co-expression of full-length HDP constructs with molecular chaperones (HSP70: DnaK-DnaJ-GrpE; HSP60: GroEL-ES) was explored to improve protein solubility. Of all the constructs tested, HDPpf showed improved solubility with HSP70 chaperones, whereas the *HDPpf-GFP* and *MBP-HDPpf* fusion constructs responded well to co-expression with HSP60 chaperonins.

#### 1.5.3.1 HDPpf with HSP70 Chaperones

Co-expression with the pKJE7 plasmid (DnaK-DnaJ-GrpE) produced a notable amount of soluble HDPpf **(Figure 2A)**. However, the protein failed to bind Ni-NTA resin under native conditions. Therefore, the soluble fraction underwent ammonium sulfate precipitation and was further processed via cation exchange chromatography (SP Sepharose, pH 5.5), where the protein eluted in the flow-through. Subsequent Ni-IDA purification under denaturing conditions (8 M urea) confirmed the identity of HDPpf **(Figure 2C–D)**. Size-exclusion chromatography (SEC) revealed two peaks **(Figure 2E)**, both containing HDPpf and DnaK, later resolved into a single monodispersed complex (**Figure 2F)** with a radius of gyration of 6.135 nm **(Figure 2G)**. Despite successful purification, the complex lacked heme-to-Hz transformation activity.

**Figure 2.**
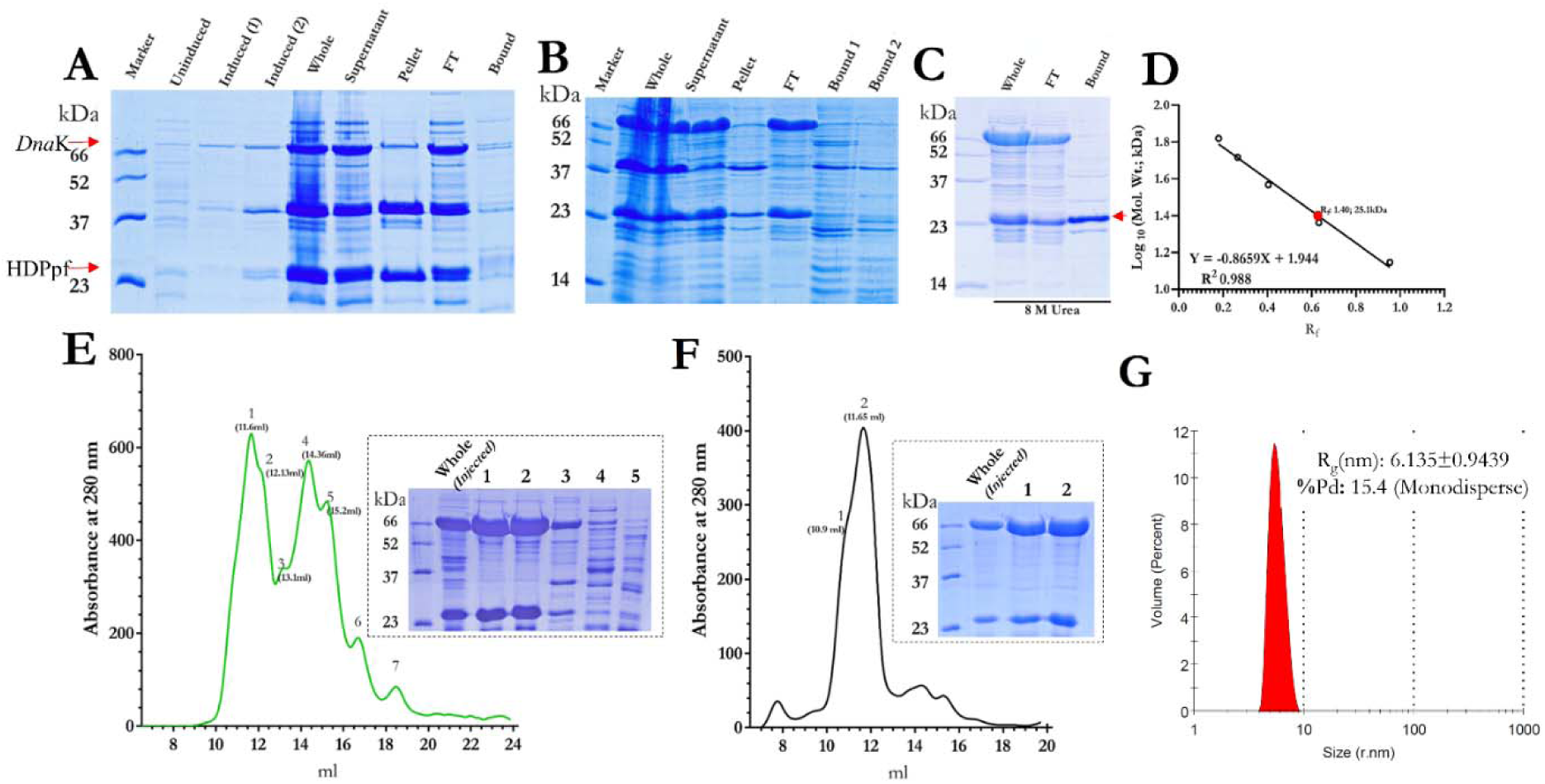
DnaK Co-expression induced solubilization of HDPpf: Coomassie Blue-stained SDS-PAGE (12% w/v) gel of various samples collected during different stages of HDPpf protein purification. **A**, Unindicted, sample exponentially growing culture before IPTG induction; Induced (1), sample of culture after L-arabinose induction; Induced (2), sample of mature culture after IPTG induction; whole, sample of lysed and homogenized slurry; supernatant, sample of supernatant obtained after centrifugation; pellet, sample of pellet obtained after centrifugation; flow-through, sample of supernatant unbound Ni-IDA beads; bound, sample bound to Ni-IDA beads. **B**, whole, sample of ammonium sulphate precipitate of flow-through sample, dissolved in CAPS buffer; supernatant, sample of supernatant obtained after centrifugation of dialyzed (AmSo4) pellet; pellet, sample of pellet obtained after centrifugation of dialized; flow-through, sample of supernatant unbound SP sepharose beads at pH5.5; bound, sample bound to SP sepharose beads. **C**, Metal immobilization chromatography of flow-through fraction (SP-sepharose chromatography) under denaturation condition of 8M urea. **D**, Molecular mass deduction of prominent band in bound lane of image C (red arrow), from Rf value (migration distance of the protein/migration distance of the dye front). The red dot corresponds to the deduced molecular mass of HDPpf. **E**, Gel-filtration purification of flow-through fraction of SP sepharose (B) chromatography, the inset is SDS-PAGE of various peaks collected during gel filtration. **F**, Gel-filtration of peak one fraction of image E, the inset is SDS-PAGE of various peaks collected during gel filtration. **G,** DLS measurement of gel-filtration fraction 2 of image E.

Small-angle X-ray scattering (SAXS) data were collected for the purified DnaK– HDPpf complex (5 mg/mL) using a lab-based Anton Paar SAXSpace instrument. Analysis was performed within the q-range of 0.011–0.4 Å⁻¹ **(Figure 3A)**. Guinier analysis (q = 0.0136–0.0324 Å⁻¹) yielded a radius of gyration (Rg) of 38.4 ± 0.11 Å (I₀ = 0.28 ± 4.46 × 10⁻_; R² = 0.997) **(Figure 3B)**. Kratky plots revealed two distinct bell-shaped curves, indicating the presence of multiple subunits within a multiprotein complex **(Figure 3C)**.

**Figure 3.**
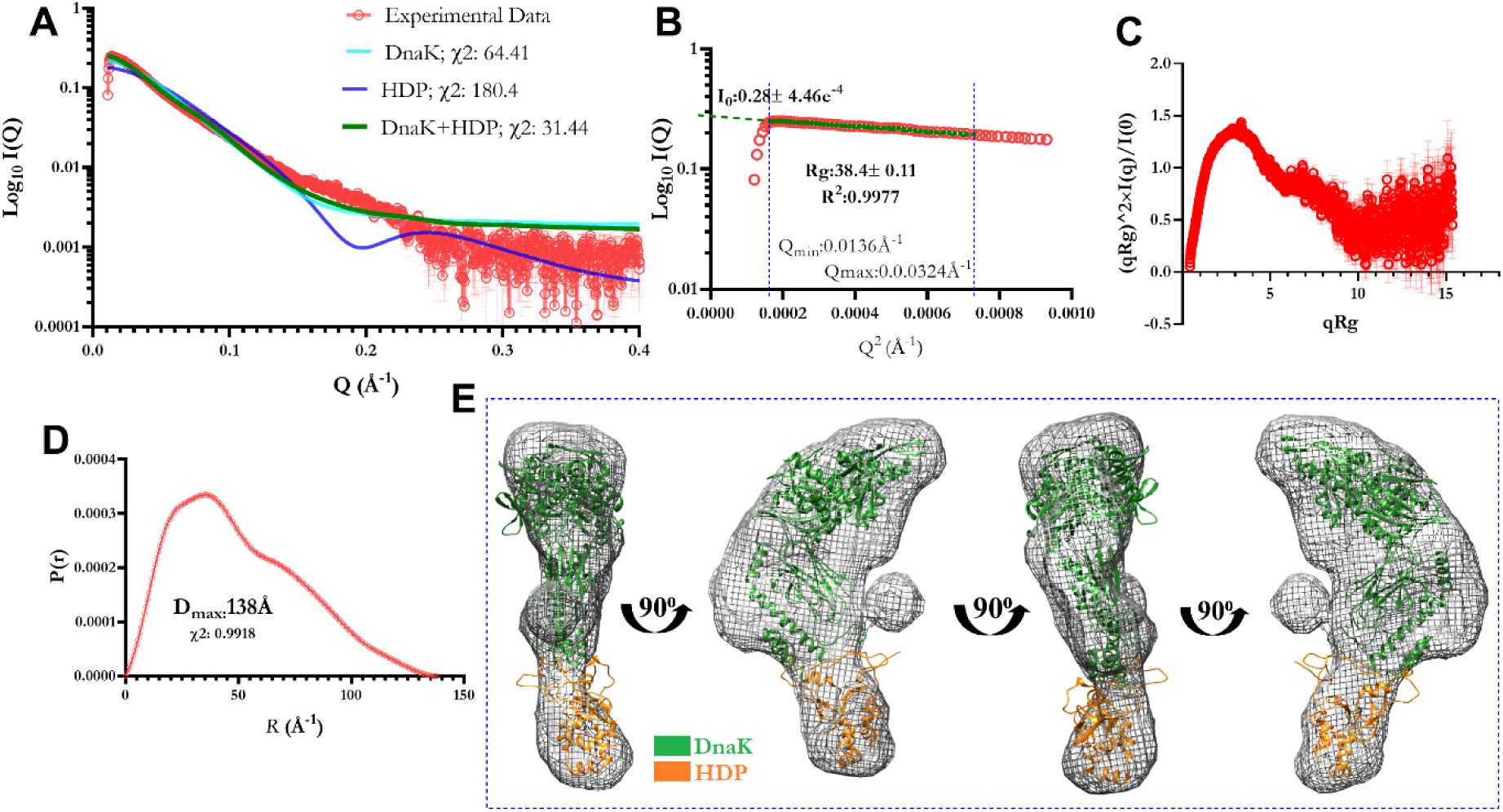
SAXS measurements on DnaK-HDPpf complex: **A.** Comparison of experimental data in red with color-coded theoretical scattering profiles of DnaK, HDP, and proposed DnaK-HDPpf complex. **B.** Guinier plot. **C.** Experimental Kratky plot. **D.** Pair distance distribution function [P(r)] for the experimental scattering data. **E.** Ab initio shape model generated with GASBOR (grey mesh; envelope at 15 Å resolution) overlaid with the crystal structure of DnaK (PDB: 7kzi; green ribbon) and HDP (predicted via AlphaFold).

Pair-distance distribution analysis (P(r)) via GNOM identified a maximum particle dimension (Dmax) of 138 Å. Ab initio modeling using GASBOR produced a 15 Å resolution envelope **(Figure 3E)**, which partially fit the crystal structure of DnaK (PDB: 7kzi). The remaining unmodeled density was successfully occupied by a predicted structure of HDPpf, supporting the formation of a DnaK–HDPpf complex. Among theoretical scattering profiles generated for DnaK, HDPpf, and their complex, the DnaK–HDPpf model best matched the experimental SAXS data (χ² = 31.44), further confirming complex formation in solution **(Figure 3A)**.

#### 1.5.3.2 HDPpf- GFP with HSP60 Chaperonins

GroEL-ES co-expression significantly enhanced the soluble expression of HDPpf- GFP. The bound fraction in Ni-IDA chromatography showed prominent bands corresponding to GroEL, HDPpf-GFP, and GFP. Gel filtration of the bound fraction indicated aggregation or a very high molecular weight of the eluted protein, except for GroEL. Anion exchange chromatography (MonoQ) at pH 9.5 resulted in the co-elution of GroEL and HDPpf-GFP, suggesting complexation **(Figure S5)**. The HDPpf- GFP fusion protein in the bound fraction was unresponsive to TEV protease digestion and did not exhibit heme-to-Hz transformation activity.

#### 1.5.3.3 MBP-HDPpf Fusion with GroEL-ES

Co-expression with GroEL-ES at 18 °C also improved solubility of the MBP-HDPpf fusion protein. Ni-IDA purification yielded two closely placed bands on SDS page **(Figure S6A)**. The upper band correspond to MBP-HDPpf (66.95 kDa) and the lower band is of GroEl (57.23 kDa) **(Figure S6B)**. along with contaminating MBP. TEV digestion partially cleaved the fusion, releasing HDPpf (∼25 kDa). SEC showed two peaks: one for free MBP and another indicating MBP-HDPpf/GroEL complexation **(Figure S6C)**. Despite the difference in their pI value (GroEl: 4.87; MBP-HDPpf: 6.67), attempts to dissociate the complex using ion exchange and hydrophobic interaction chromatography were unsuccessful, suggesting a stable interaction. No heme-to-Hz activity was observed in either the native or cleaved forms.

### 1.5.1 N-terminal Truncations Yield Soluble HDP

RoseTTAFold (PMCID: PMC7612213) structural prediction revealed that HDPpf comprises an un-structured and flexible N-terminal region (∼residues 1–50) and a structured C-terminal Fasciclin domain (residues 60–204; PFAM: pfam02469) **(Figure S7)**. While the structured domain contains nine conserved histidines essential for heme binding and transformation, the flexible-unstructured N-terminal region includes only one (His44), which is not implicated in trahsformation activity. Given the potential for enhanced solubility in globular proteins, we designed four N-terminal truncation constructs: *HDPpf-N1* (Δ11N), *N2* (Δ21N), *N3* (Δ27N), and *N4* (Δ37N), all with an N-terminal His-tag and C-terminal 3-residue truncation.

Among these, only HDPpf-N4 showed detectable soluble expression **(Figure S8A)**. Although this protein failed to bind Ni-IDA resin under native conditions, it bound under denaturing conditions (8M urea), confirming identity **(Figure S8B, S8C)**. Purification involved ammonium sulfate precipitation, followed by Q and SP ion exchange chromatography **(Figure S8D, S8E)**. Gel filtration indicated a dimeric form (∼40 kDa) **(Figure S8F).**

Although HDPpf-N4 was expressed in a soluble form, it failed to bind Ni-IDA resin and exhibited a dimeric oligomeric state during SEC analysis. This suggests that the N-terminal 6×His tag may be buried at the dimer interface, rendering it inaccessible for affinity purification. Therefore, to improve affinity purification and solubility, we redesigned ten constructs (*HDPpf-C1* to *C10*) with sequential N-terminal truncations and a C-terminal His-tag. While most constructs expressed well in Rosetta (DE3)pLysS, proteins were largely insoluble **(Figure S9A)**. Constructs *C1*, *C4*, and *C6* were only expressed in alternative hosts (C41/C43 DE3), but still formed inclusion bodies. Co-expression with DnaK/DnaJ/GrpE or GroEL/ES chaperones increased solubility for some constructs (e.g., *C3, C5, C7–C9*) but did not result in Ni-IDA-purifiable protein **(Figure S9B)**. Varying L-arabinose co-inducer levels had no effect. Most of the constructs (*C1* to *C9*), upon optimization, exhibited some expression of recombinant protein in the soluble phase **(Figure S9C)**; however, none bound to Ni-IDA resin or demonstrated measurable heme-to-Hz transformation activity.

Among all constructs, only *HDPpf-C10* (Δ44N) was reproducibly expressed in the soluble phase and retained stability **(Figure 4A)**. Homologous truncation constructs from *P. vivax* (*HDPpvC1*, Δ44N), *P. knowlesi* (*HDPpkC1*, Δ58N), *B. bigemina* (*HDPbbC1*, Δ39N), and *T. equi* (*HDPTqC1*, Δ39N) were also cloned and tested. Of these, *HDPpf-C10* and *HDPbbC1* displayed the highest expression and stability **(Figures 4B, 4C)**, enabling purification by size exclusion chromatography. *HDPpf-C10* eluted in two peaks—one at 16.66 ml (dimer, ∼44.8 kDa) and another at 19.33 ml (monomer, ∼14.8 kDa) **(Figure 4D)**. In contrast, HDPbbC1 eluted as a single monomeric peak (∼6 kDa) **(Figure 4E)**. Other constructs were either unstable or expressed solely as inclusion bodies.

**Figure 4.**
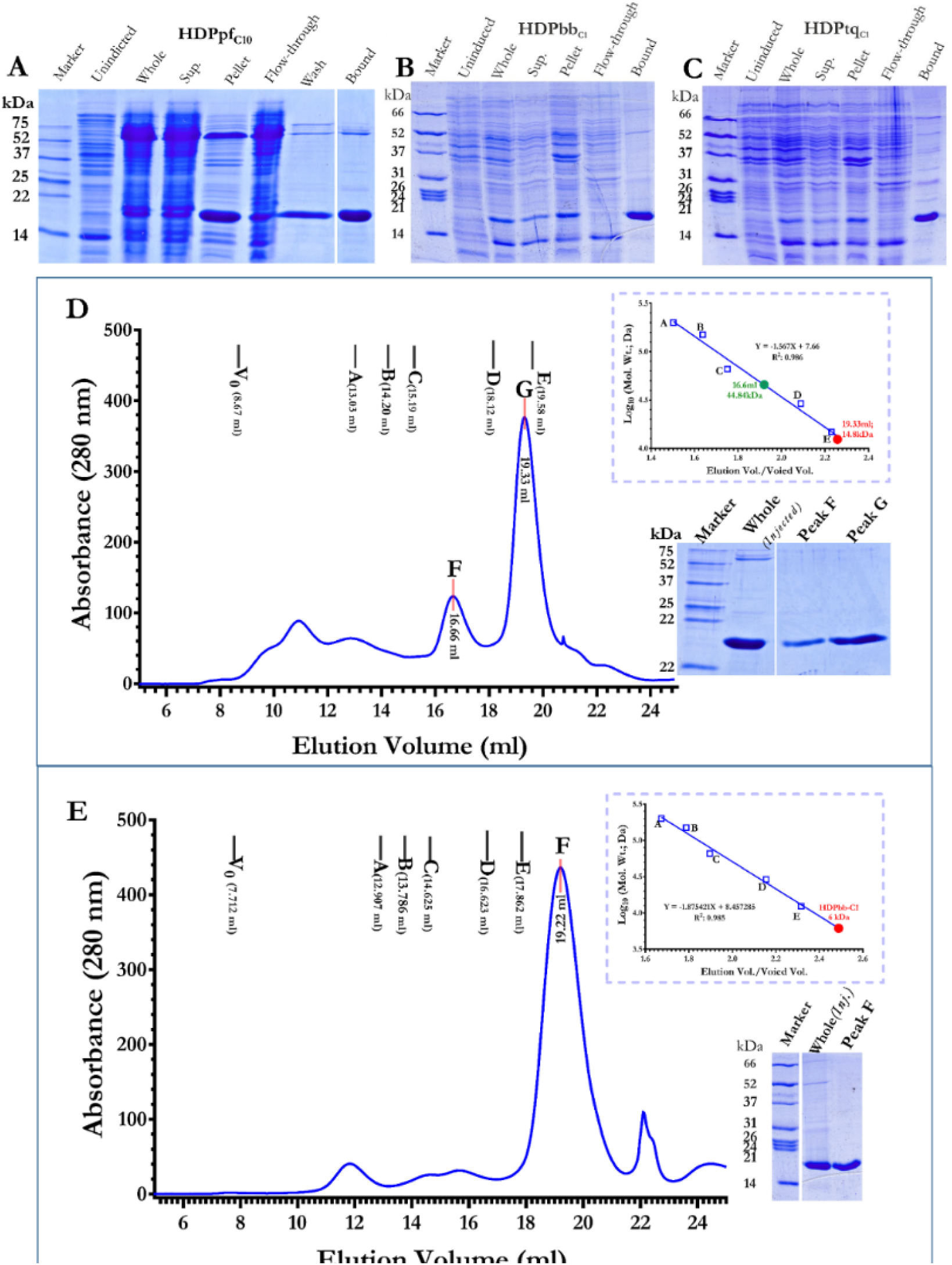
Expression and Purification of Truncated Constructs of HDP Native Soluble Phase: (**A-C**) Coomassie Blue-stained SDS-PAGE (15% w/v) gels showing the expression and purification of HDPpf-C10 (A), HDPbbC1 (B), and HDPtqC1 (C) at 18°C. The samples include: Unindicted (exponentially growing culture before IPTG induction), Whole (lysate and homogenized slurry), Supernatant (supernatant obtained after centrifugation), Pellet (pellet obtained after centrifugation), and Bound (sample bound to Ni-IDA beads). (**D-E**) Size exclusion elution profiles of HDPpf-C10 (D) and HDPbbC1 (E) purified by Ni-IDA chromatography. Molecular mass standards used for calibration are indicated: a; blue-dextran-200,000 kDa, b; β-Amylase-200 kDa, c; alcohol dehydrogenase-150 kDa, d; bovine serum albumin-66 kDa, e; carbonic anhydrase-29 kDa, f; cytochrome c-12.4 kDa. The inset shows the calibration curve for the Superdex 200 size exclusion column. The solid dot represents the deduced molecular mass of the protein corresponding to the respective peak by its elution volume. Coomassie Blue-stained SDS-PAGE (15%) gels of various samples collected during peak elution are also shown.

### 1.5.2 Soluble Constructs Lack Enzymatic Activity

Extensive efforts were made to solubilize HDP in a native-like, non-aggregated form. This form was defined as one that remains in the supernatant after centrifugation at 17,000 × g and does not elute in the void volume during size-exclusion chromatography. Multiple constructs were tested, including full-length, truncated, and fusion variants. Some of these yielded soluble recombinant proteins, either with or without binding to Ni-IDA resin. However, none of these constructs showed any measurable heme-to-Hz transformation activity.

To investigate functional integrity, we examined heme-binding properties of two representative constructs: the soluble HDPpf-C10 and full-length HDPpf solubilized from inclusion bodies using N-lauryl sarcosine **(Figure 5)**. Binding assays were conducted at pH 8 with 1 mM imidazole using spectroscopic titration, keeping heme concentration constant at 10 µM while varying protein concentrations.

**Figure. 5.**
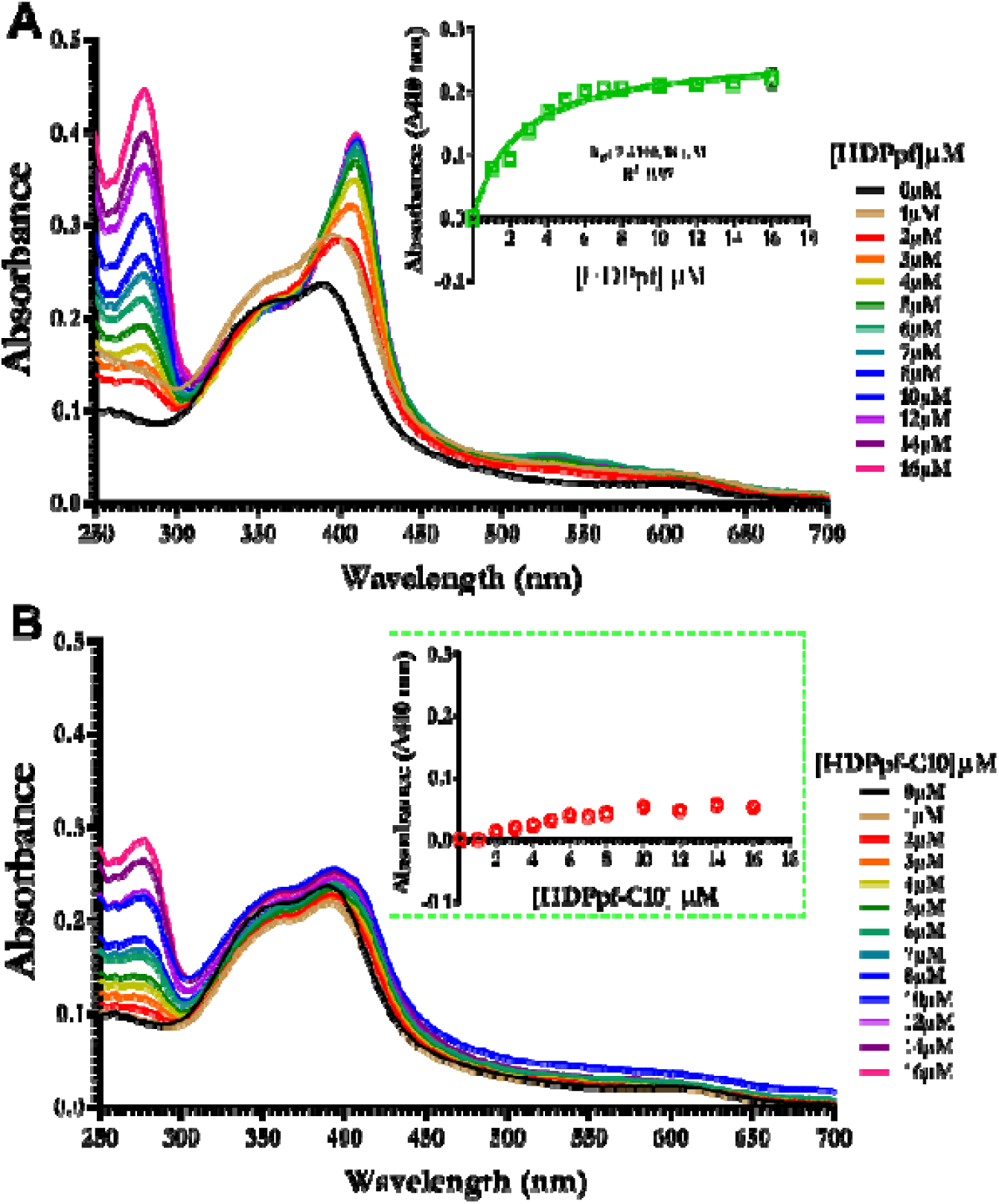
Examination of Heme Binding Characteristics of HDP Proteins via UV/VIS Spectroscopy and Titration. Panels (A) and (B) display the titration results for HDPpf and HDPpc-C10, respectively. The electronic absorption spectra represent the interaction of 10µM heme with varying concentrations of HDPpd (A) and HDPpf-C10 (B), color-coded for distinction, at pH 8 and a temperature of 37°C. The inset illustrates the change in absorbance at 410 nm (Δ410 nm=A_410_ [protein] - A_410_ [0µM protein]) relative to protein concentration, with error bars indicating the standard error across triplicate measurements.

In the absence of HDP, heme displayed a characteristic Soret band with peaks at ∼350 nm and ∼390 nm. Upon titration with full-length HDPpf, we observed a red shift of the major Soret peak to 410 nm **(Figure 5A)**, consistent with heme binding. Binding affinity analysis yielded a dissociation constant (Kd) of 2.4 ± 0.2 µM, notably higher than previously reported nanomolar-range values, possibly due to differences in experimental pH (pH 8 vs. acidic conditions in earlier studies).

In contrast, titration with HDPpf-C10 did not induce a similar spectral shift **(Figure 5B)**, indicating lack of specific heme binding. This suggests that removal of the N-terminal 44 residues—although improving solubility—may disrupt key interactions required for heme coordination, and by extension, enzymatic activity.

## 1.6 Discussion

Detoxification of host-derived heme is a critical process in the malaria parasite’s lifecycle and a validated target for antimalarial drug development. The *Plasmodium* parasite converts toxic heme, released during hemoglobin digestion, into inert crystalline hemozoin (Hz) within its acidic food vacuole [17]. The heme detoxification protein (HDP) is the most efficient known biological mediator of this transformation. Despite its central role, mechanistic and structural studies of HDP have been hampered by the inability to express it recombinantly in a soluble, detergent free native form.

In this study, we systematically explored multiple strategies to achieve native, soluble expression of HDP in *E. coli*. These approaches included expression of orthologs, consensus sequence design, fusion to solubility-enhancing partners (e.g., MBP, TRX, GFP, pCold-TF), co-expression with molecular chaperones (DnaK/DnaJ/GrpE and GroEL/ES), and extensive construct optimization through N-terminal truncations. We successfully expressed full-length HDPpf in a soluble form by co-expressing it with the DnaK/DnaJ/GrpE chaperone system. However, the recombinant protein failed to bind Ni-IDA resin, indicating it was likely in complex with DnaK due to a misfolded or incompletely folded state, and showed no detectable heme-to-Hz transformation activity. Similar outcomes were observed with HDPpf-GFP and MBP-HDPpf fusion constructs co-expressed with GroEL/ES, where soluble protein was obtained but formed stable complexes with the chaperonin, again suggesting misfolding and resulting in enzymatic inactivity. In the case of the pCold-HDPpf fusion protein, protease cleavage of the soluble fusion led to the precipitation of HDPpf into the pellet fraction, further confirming improper folding. These repeated observations support the idea that full-length HDP may only attain a stable tertiary structure in the context of its native hetero-oligomeric quaternary assembly, as suggested by earlier studies [17]

Upon bioinformatic analysis **(Figure S10)**, the N-terminal region of HDP was predicted to be intrinsically disordered and potentially involved in protein–protein interactions. Interestingly, HDP contains nine conserved histidine residues, of which five were previously implicated in heme binding and transformation activity [28] [29]. Early studies identified His122, His172, His175, and His197 as key residues for heme binding [28], but subsequent mutagenesis and structural investigations revised this understanding, confirming the role of His172, His175, His192, and His197, while discounting His122 [29]. Notably, our analysis suggests that while the structured FAS1 domain harbors most of these critical histidines, the N-terminal ∼50 residues are unstructured/ flexible and include only one histidine, His44, which has been shown not to participate in heme binding.

Given that expression of globular, well-folded proteins is generally more successful in *E. coli*, and considering the likely destabilizing influence of the flexible-unstructured N-terminal region, we rationalized that truncating this segment could improve expression without compromising functionality. This rationale led to a series of N-terminal truncation constructs. Among them, the *HDPpf-C10* construct, lacking the first 44 residues but retaining all functionally characterized histidines, was expressed in a soluble form and readily purified. However, it failed to demonstrate either heme-binding or heme-to-Hz transformation activity. These findings suggest that the flexible unstructured N-terminal region, despite its lack of direct involvement in heme coordination, plays an indispensable role in proper protein folding and functional integrity.

This region may be required to stabilize the overall tertiary structure or to mediate essential interactions with other food vacuole proteins such as falcipain-2, plasmepsin-2/4, and histo-aspartic proteases—partners with which HDP is known to form native complexes [17]. Such stabilization is likely absent in the bacterial expression system, but may be partially mimicked by detergents, which have been shown to preserve HDP activity in solubilized inclusion body preparations. Thus, our findings underscore the functional importance of intrinsically disordered segments and the need for expression systems that better replicate the native cellular context of *Plasmodium*.

A key limitation of our study is its confinement to *E. coli*-based expression systems. While this platform offers scalability and ease of manipulation, it lacks the cellular machinery required to fold complex or membrane-associated eukaryotic proteins correctly. Alternative systems, such as yeast, insect cells, or cell-free eukaryotic systems, may offer a more suitable environment for native folding and function. Nevertheless, our comprehensive *E. coli*-based screen provides a critical framework for construct design, solubility profiling, and purification workflows that will aid future expression efforts in more advanced systems.

In conclusion, our findings highlight the dual challenge of HDP expression: improving solubility without compromising function. While N-terminal truncations enhance solubility, they abolish Heme-to-Hz transformation activity—underscoring the essential, non-redundant role of the flexible-unstructured N-terminal domain. Future efforts should focus on co-expression with native binding partners or transition to eukaryotic hosts to fully reconstitute HDP’s native structure and catalytic mechanism.

## Supporting information

Supplementary information comprising commercially synthesized coding dna and pcr primer sequences and ten supplemental figures, labeled S1 to S10.

## 1.7 Acknowledgement

We wish to express our sincere appreciation to Mr. Sitanshu Kumar Sarangi, an M.Sc. Biochemistry student from the Department of Biotechnology at Ravenshaw University, Cuttack (Trainee; January to June 2017), and Ms. Monika Pandey, a student from the School of Life Sciences at Devi Ahilya Vishwavidyalaya, Indore (Trainee; January to June 2019). Both individuals were exposed to recombinant DNA technology under our supervision, and they made significant contributions to our research on the cloning and expression of *Plasmodial* genes.

## 1.8 Abbreviations

HDP: Heme Detoxification Protein
Hz: Hemozoin / β-Hematin
CDS: Coding DNA Sequence
STR: Streptavidin tag
TEV: Tobacco Etch Virus (protease cleavage site)
TRX: Thioredoxin
MBP: Maltose Binding Protein
GFP: Green Fluorescent Protein
TF: Trigger Factor
IMAC: Immobilized Metal Affinity Chromatography
Ni-IDA: Nickel–Iminodiacetic Acid
IPTG: Isopropyl β-D-1-thiogalactopyranoside
SAXS: Small-Angle X-ray Scattering
Rg: Radius of Gyration
I(0): Scattering Intensity at q = 0
P(r): Pair Distance Distribution Function
SEC: Size-Exclusion Chromatography
DLS: Dynamic Light Scattering
CAPS: N-cyclohexyl-3-aminopropanesulfonic acid
SDS-PAGE: Sodium Dodecyl Sulfate–Polyacrylamide Gel Electrophoresis
HSP: Heat Shock Protein
DnaK/J/GrpE: Components of the HSP70 chaperone system
GroEL/ES: HSP60 chaperonin complex
pI: Isoelectric Point
q: Scattering vector magnitude in SAXS
χ²: Chi-squared statistical measure of fit in SAXS analysis
PCR: Polymerase Chain Reaction
OD: Optical Density (at 600 nm)
BLAST: Basic Local Alignment Search Tool
UniProt: Universal Protein Resource.

## 1.9 Data availability

All data are contained within the manuscript.

## 1.10 Supporting information

The manuscript includes supplementary information comprising commercially synthesized coding dna and pcr primer sequences and ten supplemental figures, labeled S1 to S10.

## 1.11 Author’s contribution

RDM and RS nucleated the idea; RS performed all experiment and analysis; RS and RDM wrote the manuscript.

