## Supplementary information comprising commercially synthesized coding dna and pcr primer sequences and ten supplemental figures, labeled S1 to S10. for "Recombinant Expression and Purification of *Plasmodium* Heme Detoxification Protein in *E. coli:* Challenges and Discoveries"

### *Suplimentary information*

#### Suplimentary information content:

1. **Commercially synthesized Coding DNA and PCR primer Sequences**

A. Heme detoxification protein from *Plasmodium falciparum;* UniProt: Q8IL04

*>HDPpf, Plasmodium falciparum; UniProt: Q8IL04*

cgtggatccAAGAACCGTTTTTATTACAACTTGATCATTAAGCGTCTTTACACCCGTT
CCGGTGGTTTACGTAAACCCCAGAAGGTTACTAACGATCCCGAAAGTATCAACC
GTAAGGTCTACTGGTGCTTCGAACATAAACCTGTCAAACGTACCATTATTAACTT
GATCTACTCGCATAACGAACTTAAAATTTTTTCGAATCTTTTGAATCACCCTACG
GTAGGATCGTCTTTGATTCACGAGCTGTCATTGGACGGCCCATACACCGCTTTT
TTCCCTAGTAATGAGGCCATGCAGCTTATTAACATCGAGAGCTTCAACAAATTGT
ATAACGATGAGAATAAACTGTCAGAGTTTGTGTTGAATCACGTAACTAAAGAGTA
CTGGCTTTATCGTGATTTATATGGATCGAGCTACCAGCCCTGGTTAATGTATAAT
GAGAAGCGTGAAGCACCAGAAAAACTGCGTAACCTGTTGAATAATGATCTGATC
GTGAAAATCGAAGGTGAGTTCAAACATTGCAACCATTCGATTTATCTTAATGGCT
CGAAGATTATCCGTCCGAACATGAAATGCCATAATGGTGTCGTGCACATCGTCG
ATAAACCAATCATCTTTTAAgcttacg

B. Heme detoxification protein from Plasmodium Vivax; UniProt: A0A1G4HHQ7

*>HDPpv, Plasmodium Vivax; UniProt: A0A1G4HHQ7*

cgtggatccAAAAAATCCCGTCCACCTTTTTTAGTTATTAAGCGTCTGTATACTCGTT
CGGGAGGCTTACGTAAACCGCAAAAGGTCACGAACGATCCGGAAAGTATCAAC
CGTAAAACTTATTGGTGTTTTGAGCATAAGCCAATCAAGCGTACCTTAGTCAACT
TGATTTACAGCCATAATGAGCTTAAATTGTTTAGCCGTTTTTTGAATCACCCAAA
CGTGGGTACTTCCCTTGTGCATGAATTGTCCTTAGAAGGGCCATATACAGGATT
CTTACCCAGTAATGAAGCCTTGAAACTTATTTCGCCGGAGAGCTTGGCTAAGTT
GTACGAAGAAGGGGACAAGCTGATGGAGTTTGTCTTGGGCCATTTTGCCAAAG
ACTTCTGGCTGTATCGTGACTTATATGGCTCATCCTATCAACCCTGGTTAGTGTT
TAACGAGCGTCGTGACGCCCCAGAAAAGATTACCAATCTGGTCAATCGTGATCT
GTTAGTCGAAATCACTGGAGAGTTCAAAAATTGTGACCACTCTATTTCTCTTAAT
GGTGCTAAGATTATCCGTCCCAATATGAAATGCCATAACGGTGTCGTACACATT
GTGGACCGTCCAATTATCCAACGTTAAgcttacg

C. Heme detoxification protein from *Babesia bigemina;* UniProt: A0A061DBU8

*> HDPbb, Babesia bigemina; UniProt: A0A061DBU8*

cgtggatccAGCAGCCGTCGTGCATTACAGTGGGGACCCCGTCCAATCAATGCTGG
TCGTTTACGTCGTGTAACCCGTAACCCCGACGTTATTAACCGTTTGGTGTACTGG
TGTTTCGAGCATCCAGAAGTTCGTCGTACTGTGCTTAACGCATGCCAGGGAGGT
AAATTCACTCGTTTATTCGGTTCCCTTTTCAATGTGGAAACCGCACACGGGTATG
GGCTTGCACACGAATTATCTCTGCCGGGGCCTTTCACCGGCTTCATTCCGGTGG
ATGACGGACTTCAAAAGTTAATGCACACGTTATATGAATCTCCACCTGAACGTATC
GCAGACTTTGTTCGTTCACACTTCACCCGTGACTTATGGCTTCACCGTGACATCA
CAGGCTCGCCTATGCAACCTTGGCTGCCCTACAACGCTCCACGTGCAGCACCT
TCGAGTTTATTGGCTCTTTCGGGGCGTGAGTTAGCCGTTGAAAATGGAGGAGAC
GTTCGTATTAAGACAGATTATACAACCATCAACGGGAGTAAAGTTTTACGTTGGAA
CATGCGTTGCCATAATGGTGTCATTCATTTAGTGGATCGTCCGGTGATTTTAGAA
GACCTTTAAgcttacg

### *Suplimentary information*

D. Heme detoxification protein from *Theileria equi;* UniProt: L0B1I2

*>HDPtq, Theileria equi; UniProt: L0B1I2*

cgtggatccCACCTGTCAAGTCACCGTAACTGGCGTAGTACCTTTGGCGGTTTGAAA
AAAGTCCGTGGCACGACGAAAGATCCAGAGGTTGTCAATCACAAGGTATATTGG
TTCTCAGAGCATCGTAATGTTTGCCGTACTGTATTGAGTTTATGTCGTACACATCC
GTTTACAAACATCTTTTCCGCACTTATCGACCCTGAGACCACCTCAGGGTATGCC
ATTAGTCACGAGCTGTCGTTGCCTGGCCGTTTACCGGGTTTATCCCCGTAGAC
AACGGAATGAAACAGATCATCAAGCAAATTGAGCGTAAAGGTAACGATTTTCTTG
TTGATTTCATTCGTTCCCACTTCACTTTAGACTTATGGTTACATCGTGACATTACTG
GCTCGTCCACCCAACCTTGGTTGTTATATAATAAGGAGCGTAAGGCCCCTGAACA
CTTACTTTCGCTTACCAACTCTAAGCTGGTTATTGAGAATATCGGAGAAATCTCAC
GTGGTACAGACTTAACTCATATTAATGGTTCTAAAATTCTTCGTTGGAACATGCGT
TGCCACAATGGGGTCATTCATCTGATCGACCGTCCCATGATGGGGCTGTAAgctta
cg

E. Heme detoxification protein from *Plasmodium knowlesi;* UniProt: A0A384L7P6

> HDPpk, *Plasmodium knowlesi;* UniProt: A0A384L7P6

cgtGGATCCCTGATTATCAAGCGTTTGTATACTCGTTCAGGCGGCCTTCGTAAACC
GCAGAAGGTGACTAACGATCCTGAGTCGATCAATCGTAAAACATATTGGTGTTTT
GAACACAAACCCATCAAGCGTACGATGGTGAATTTGATTTACTCACACAATGAGT
TGAAATTATTCTCTCGTTTTCTTAGCCATCCTAACGTGGGAACTTCTTTGATCCAC
GAGCTGTCGTTGGAGGGGCCTTACACAGGATTCCTGCCTTCAAATGAGGCCTT
GAAACTTATTTCACCAGAGTCCCTTGCCAAATTATACGAACAGGGAGACAAGTTA
ATGGAATTTGTATTAGGCCACTTTACGAAGGATTTCTGGTTGTATCGTGATCTGTA
CGGTTCATCATACCAGCCATGGTTGGTATTTAATGAGAAACGTGAAGCCCCGGAA
AAAATTACCAATTTAGTCAACAAGGACCTTTTGGTCAAAATCACTGGTGAGTTCA
AAAATTGCGATCATAGTATCTTTTTGAACGGAGCCAAGATTATTCGTCCCAATATG
AAATGTCACAACGGCGTCGTTCACATTGTAGACCGTCCGATCTAAtgtacaacg

F. Heme detoxification protein with consensus design

*>HDPcc, Consensus design*

cgtggatccATGAAAAAAAAACTGTACTTCCTGGTTATCAAACGTCTCTACACCCGTT
CTGGTGGTCTGCGTAAACCGCAGAAAGTTACCAACGACCCGGAATCTATCAACC
GTAAAGTTTACTGGTGCTTCGAACACAAACCGATCAAACGTACCGTTGTTAACCT
GATCTACTCTCACAACGAACTGAAAATCTTCTCTAACCTGCTGAACAACCCGAAC
GTTGGTTCTTCTCTGATCCACGAACTGTCTCTAGAAGGTCCGTACACCGCGTTC
CTGCCGTCTAACGAAGCGCTGAACCTGATCAACATCGAATCTCTGAACAAACTG
TACGAAGACGACAACAAACTGATGGAATTCGTTCTGAACCACGTTACCAAAGAC
CTGTGGCTGTACCGTGACCTGTACGGTTCTTCTTACCAGCCGTGGCTGGTTTAC
AACGAAAAACGTGAAGCGCCGGAAAAAATCCGTAACCTGGTTAACAACGACATC
ATCGTTAAAATCGAAGGTGAATTCAAAAACTGCGACCACTCTATCTACCTGAACG
AAGCGAAAATCATCCGTCCGAACATGAAATGCCACAACGGTGTTGTTCACATCG
TTGACAAACCGATCATCTTCCAGtgtacaacg

Annexure Table *S1*: List of used PCR Primers

| Sr. No. | Construct Name | Restriction site | Direction | Sequence* | |
|---|---|---|---|---|---|
| 1. | HDPpf-N1 | BamH1 | Forward | cgtggatccAAGCGTCTTTACACCCGTTC | Commercially synthesizes CDS of HDP from *Plasmodium falciparum* |
| | | Hind3 | Reverse | cgtaagcttaTTTATCGACGATGTGCACGAC | |
| 2. | HDPpf-N2 | BamH1 | Forward | cgtggatccTTACGTAAACCCCAGAAGGTTA | |
| | | Hind3 | Reverse | cgtaagcttaTTTATCGACGATGTGCACGAC | |
| 3. | HDPpf-N3 | BamH1 | Forward | cgtggatccACTAACGATCCCGAAAGTATCAA | |
| | | Hind3 | Reverse | cgtaagcttaTTTATCGACGATGTGCACGAC | |
| 4. | HDPpf-N4 | BamH1 | Forward | cgtggatccGTCTACTGGTGCTTCGAACATA | |
| | | Hind3 | Reverse | cgtaagcttaTTTATCGACGATGTGCACGAC | |

| | | | | |
|---|---|---|---|---|
| **5.** | HDPpf-C1 | Nco1 | Forward | catgccatgggcAAGAACCGTTTTTATTACAACTTGATCATT |
| | | Xho1 | Reverse | cagtctcgagAAAGATGATTGGTTTATCGACGATGTG |
| **6.** | HDPpf-C2 | Nco1 | Forward | catgccatgggcTTGATCATTAAGCGTCTTTACACCC |
| | | Xho1 | Reverse | cagtctcgagAAAGATGATTGGTTTATCGACGATGTG |
| **7.** | HDPpf-C3 | Nco1 | Forward | catgccatgggcGGTTTACGTAAACCCCAGAAGG |
| | | Xho1 | Reverse | cagtctcgagAAAGATGATTGGTTTATCGACGATGTG |
| **8.** | HDPpf-C4 | Nco1 | Forward | catgccatgggcAAACCCCAGAAGGTTACTAACGATC |
| | | Xho1 | Reverse | cagtctcgagAAAGATGATTGGTTTATCGACGATGTG |
| **9.** | HDPpf-C5 | Nco1 | Forward | catgccatgggcAAGGTTACTAACGATCCCGAAAGTA |
| | | Xho1 | Reverse | cagtctcgagAAAGATGATTGGTTTATCGACGATGTG |
| **10.** | HDPpf-C6 | Nco1 | Forward | catgccatgggcAACGATCCCGAAAGTATCAACCGT |
| | | Xho1 | Reverse | cagtctcgagAAAGATGATTGGTTTATCGACGATGTG |
| **11.** | HDPpf-C7 | Nco1 | Forward | catgccatgggcGAAAGTATCAACCGTAAGGTCTACT |
| | | Xho1 | Reverse | cagtctcgagAAAGATGATTGGTTTATCGACGATGTG |
| **12.** | HDPpf-C8 | Nco1 | Forward | catgccatgggcAACCGTAAGGTCTACTGGTGCTT |
| | | Xho1 | Reverse | cagtctcgagAAAGATGATTGGTTTATCGACGATGTG |
| **13.** | HDPpf-C9 | Nco1 | Forward | catgccatgggcGTCTACTGGTGCTTCGAACATAAAC |

| | | | | | |
|---|---|---|---|---|---|
| | | Xho1 | Revers e | cagtctcgagAAAGATGATTGGTTTATCGACG ATGTG | |
| **14.** | HDPpf-C10 | Nco1 | Forward | catgccatgggcAAACCTGTCAAACGTACCATT ATTAACTT | |
| | | Xho1 | Revers e | cagtctcgagAAAGATGATTGGTTTATCGACG ATGTG | |
| **15.** | HDPtq-C1 | Nco1 | Forward | catgccatgggcCGTAATGTTTGCCGTACTGTA TTGA | HDPtq |
| | | Xho1 | Revers e | cagtctcgagCAGCCCCATCATGGGACGGT | |
| **16.** | HDPpk-C1 | Nco1 | Forward | cgttctagatgAAACCCATCAAGCGTACGATGG T | HDPpk |
| | | Xho1 | Revers e | cagtctcgagGATCGGACGGTCTACAATGTGA A | |
| **17.** | HDPbb-C1 | Nco1 | Forward | catgccatgggcCCAGAAGTTCGTCGTACTGT GC | HDPbb |
| | | Xho1 | Revers e | cagtctcgagAAGGTCTTCTAAAATCACCGGA CG | |
| **18.** | HDPpv-C1 | Nco1 | Forward | catgccatgggcAAGCCAATCAAGCGTACCTTA GTC | HDPpv |
| | | Xho1 | Revers e | cagtctcgagACGTTGGATAATTGGACGGTCC A | |

** Lowercase sequences do not match the template DNA, as they primarily contain sequences for restriction enzyme sites, stop codons, and additional bases necessary for the efficient functioning of restriction enzymes*

### *Suplimentary information*

#### 2. Supplimentary Figure S1 to S10

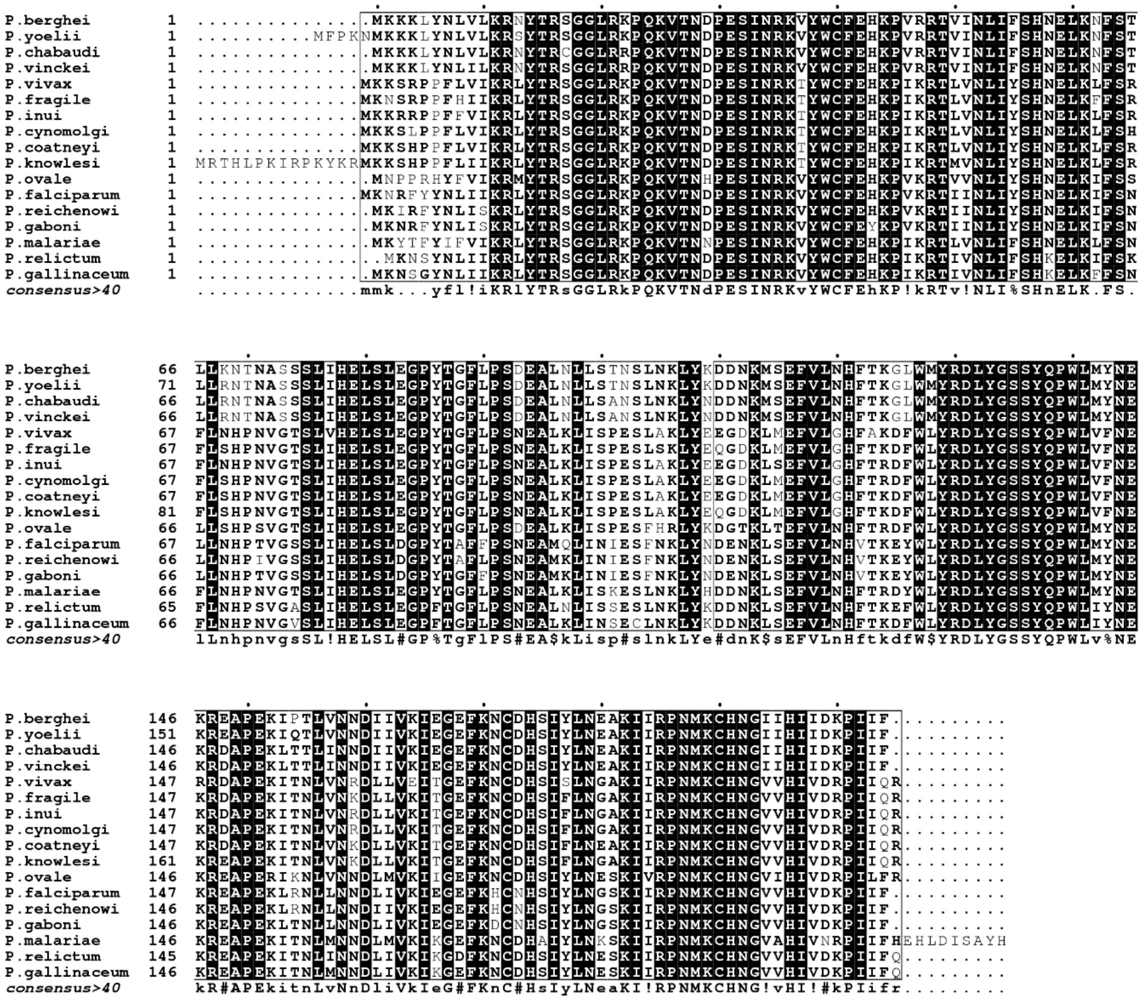

Figure S1. Multiple sequence alignment of 17 homologues of HDP from 17 different species of *Plasmodium* used to make consensus design construct of HDP (image prepared by using "ESPript3" online tool)

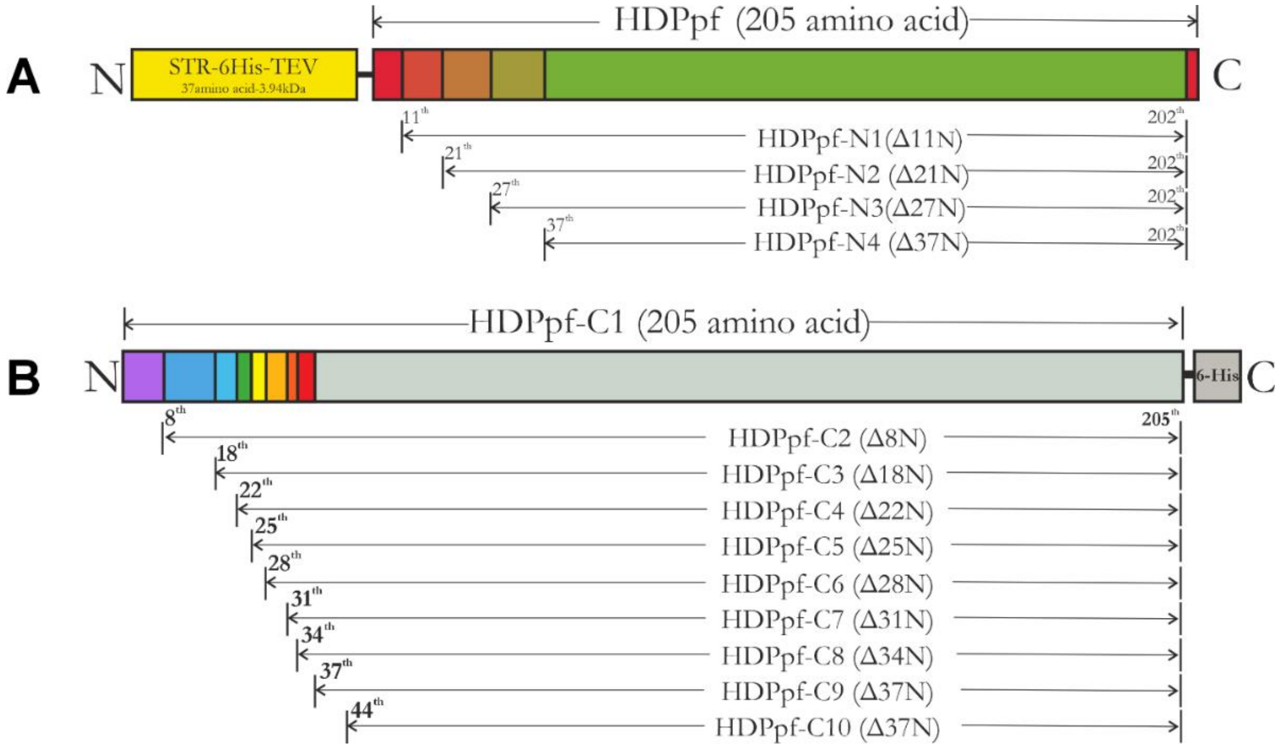

Figure S2. Schematic representation of various cloned HDPpf constructs with N-terminal truncations, featuring **(A)** N-terminal STR-6His-TEV tag, and **(C)** C-terminal 6-His tag.

### *Suplimentary information*

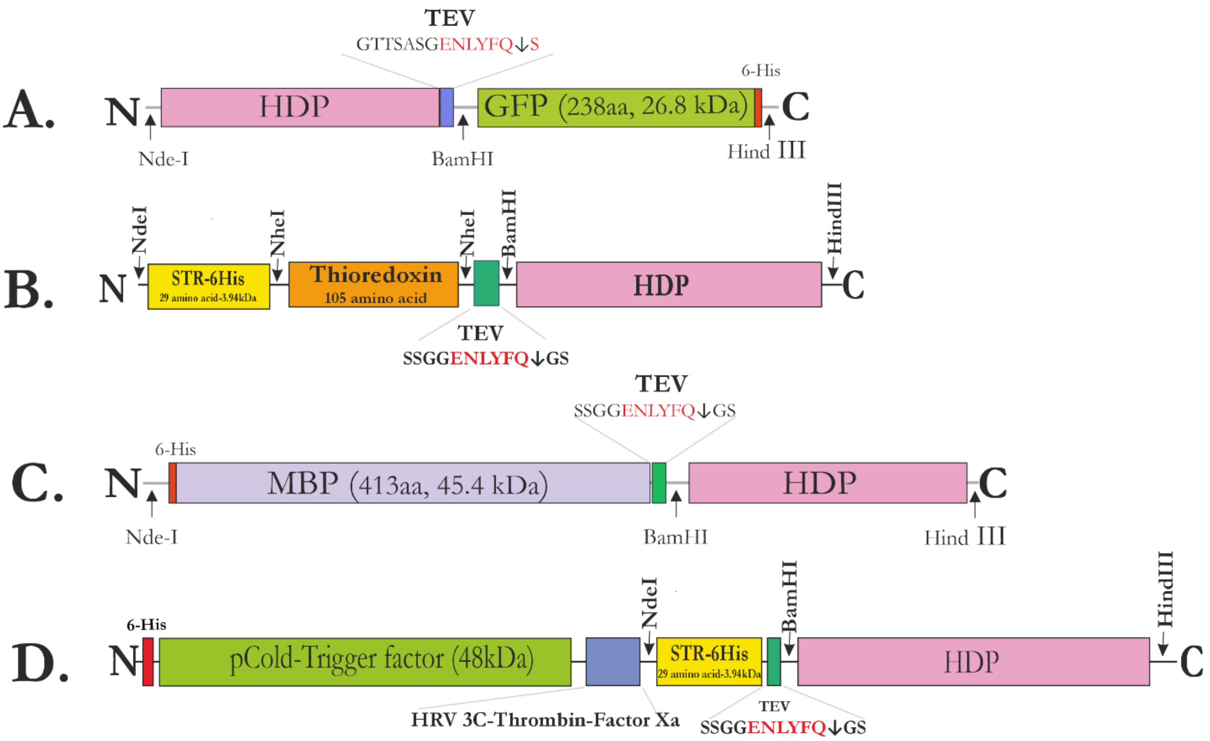

Figure S3. Schematic representation of fusion protein construct. **A**, HDP-GFP fusion protein construct. **B,** TRX-HDP fusion protein construct. **C**, MBP-HDP fusion protein construct. **D**, pCold-HDP fusion protein construct.

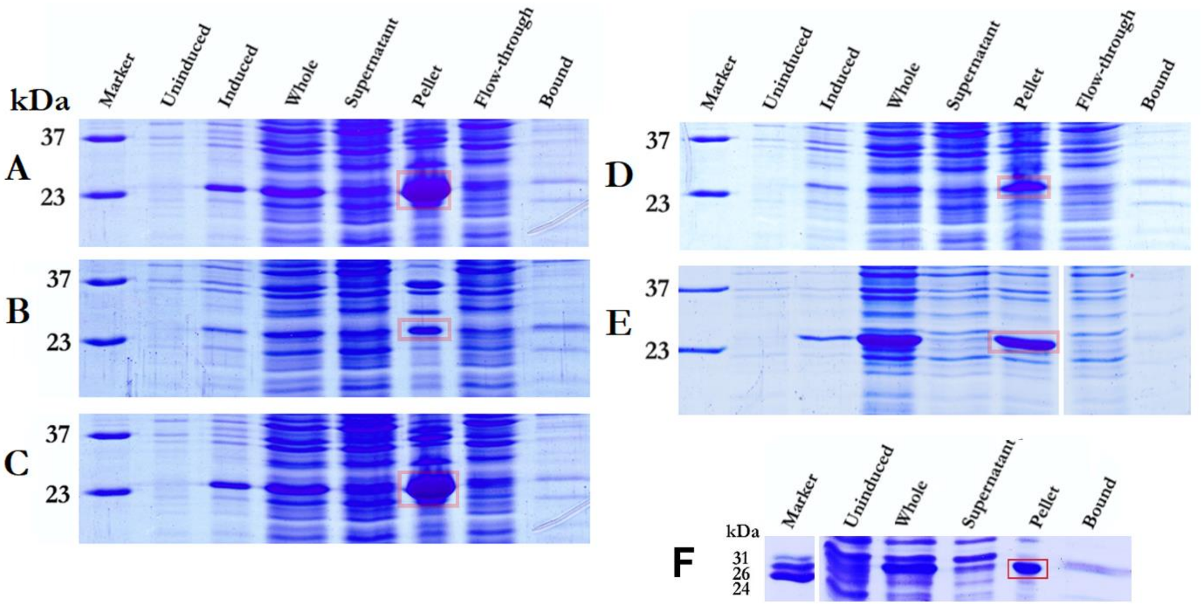

Figure S4. Expression of HDP constructs: Coomassie Blue-stained SDS-PAGE gel of several samples collected during various stages during expression and purification of **(A)** HDPpf, **(B)** HDPpv, **(C)** HFPpk **(D)** HDPtq **(E)** HDPbb **(F)** HDPcc at 18˚C. Unindicted, sample of exponentially growing culture before IPTG induction; Induced, sample of mature culture after IPTG induction; whole, sample of lysed and homogenized slurry; supernatant, sample of supernatant obtained after centrifugation; pellet, sample of pellet obtained after centrifugation; flow-through, sample of supernatant unbound to Ni-IDA resin; bound, sample bound to Ni-IDA resin. The band corresponding to HDP proteins is highlighted by red box

### *Suplimentary information*

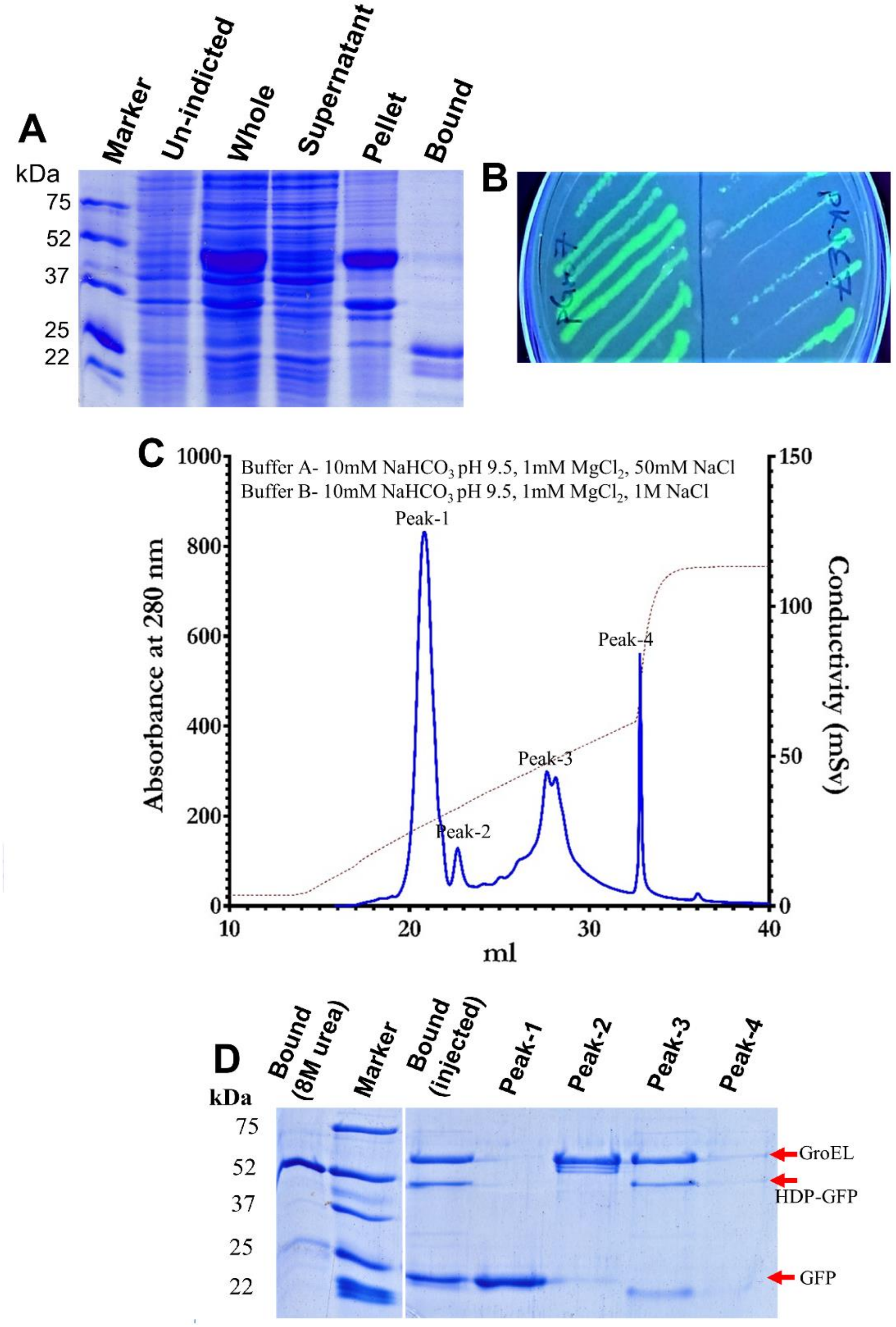

Figure S5. Expression of HDPpf-GFP fusion protein: **A**, Coomassie Blue-stained SDS-PAGE gel (15% W/V) displaying various samples collected during the expression of HDPpf-GFP fusion protein at 18°C. **B**, C41(DE3) cells expressing HDPpf- GFP fusion protein with Co expression of GroEl (left) and DnaK (Right) on LB plate with IPTG and L-arabinose. **C**. Elution profiles of MonoQ anion chromatography for the bound fraction purified during expression of HDPpf- GFP construct along with GroEl-ES. Absorbance at 280nm is shown on the left y-axis, and conductivity is shown on the right y-axis. **D,** The Coomassie Blue-stained SDS-PAGE gel (12% W/V) on the right displays various samples collected during protein elution. The "Bound (8M urea)" represents the bound fraction of Ni-IDA chromatography with inclusion bodies dissolved in 8M urea, while "Bound (injected)" corresponds to the bound fraction eluted during native co-expression of HDPpf-GFP fusion protein along with pGro7.

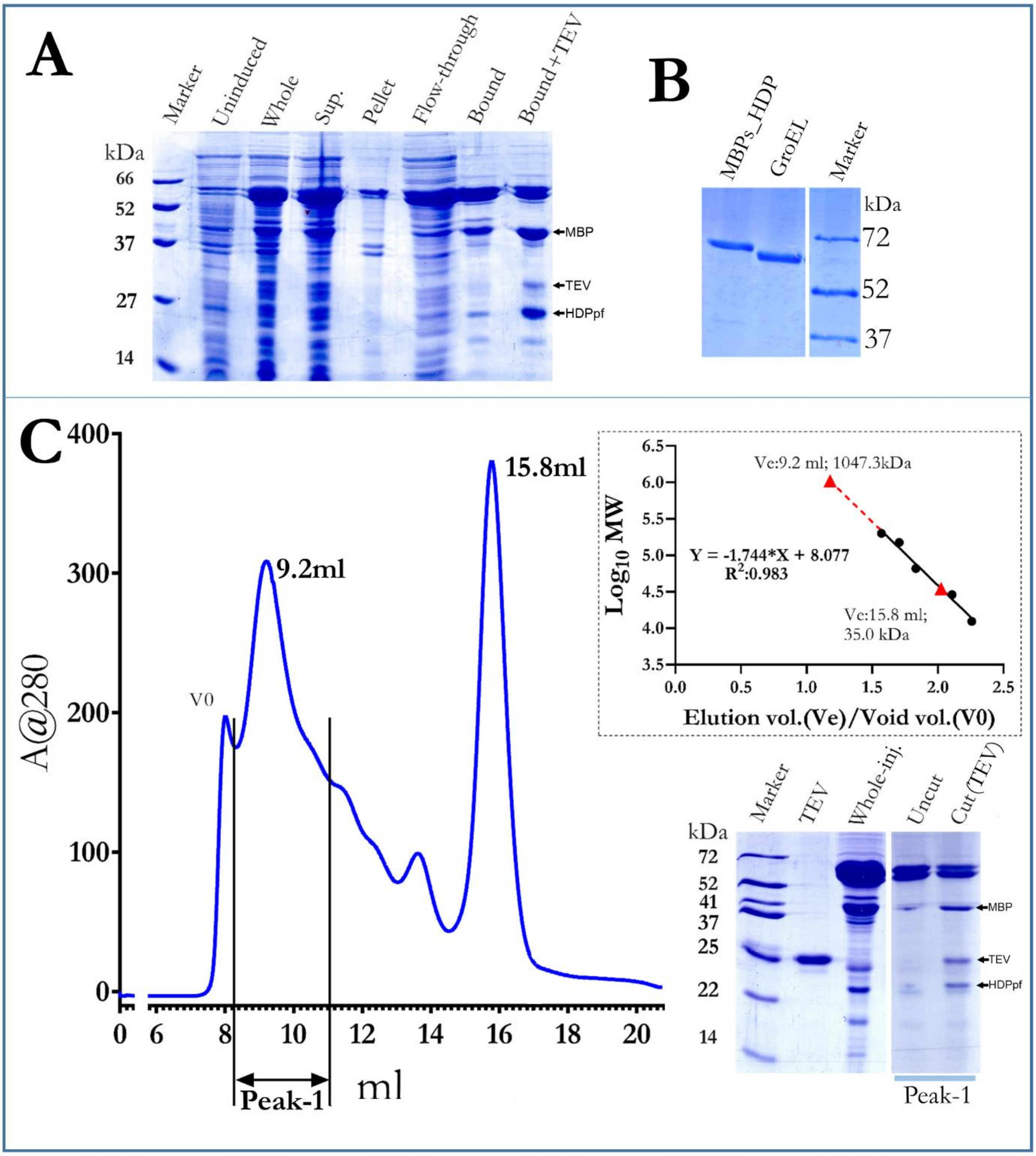

Figure S6. Co-expression of MBP-HDPpf fusion protein construct along with pGro7. **A,** Coomassie Blue-stained SDS-PAGE (12% W/V) gel of various fractions collected during co expression of pGro7 along with MBP-HDPpf fusion construct at 18ºC. Uninduced, sample exponentially growing culture before IPTG induction; Whole, sample of lysed and homogenized slurry after IPTG induction; supernatant, sample of supernatant obtained after centrifugation; pellet, sample of pellet obtained after centrifugation; flow-through, sample of supernatant unbound Ni-IDA beads; bound, sample bound to Ni-IDA beads; Bound+TEV, bound sample digested with TEV peptidase. **B**; Coomassie Blue-stained SDS-PAGE (12% W/V W/V) gel of MBP-HDPpf fusion protein purified via Ni-NTA under denaturating condition of 8M urea and purified GroEl. **C**; Size exclusion chromatography elution profile of recombinant MBP-HDPpf fusion protein co-expressed with GroEL. The inset shows the calibration curve for the Superdex 200 size exclusion column. The solid triangular dot represents the deduced molecular mass of the protein corresponding to the respective peak by its elution volume. Coomassie Blue-stained SDS-PAGE (12%) gel of various fractions collected during gel-filtration of MBP-HDPpf fusion protein. TEV; Tev peptidase, Whole-inj.; the protein injected into the columnUncut; Gel-filtration fraction (peak-1) not digested with Tev peptidase, Cut; Gel-filtration fraction (peak-1) digested with TEV peptidase.

#### *Suplimentary information*

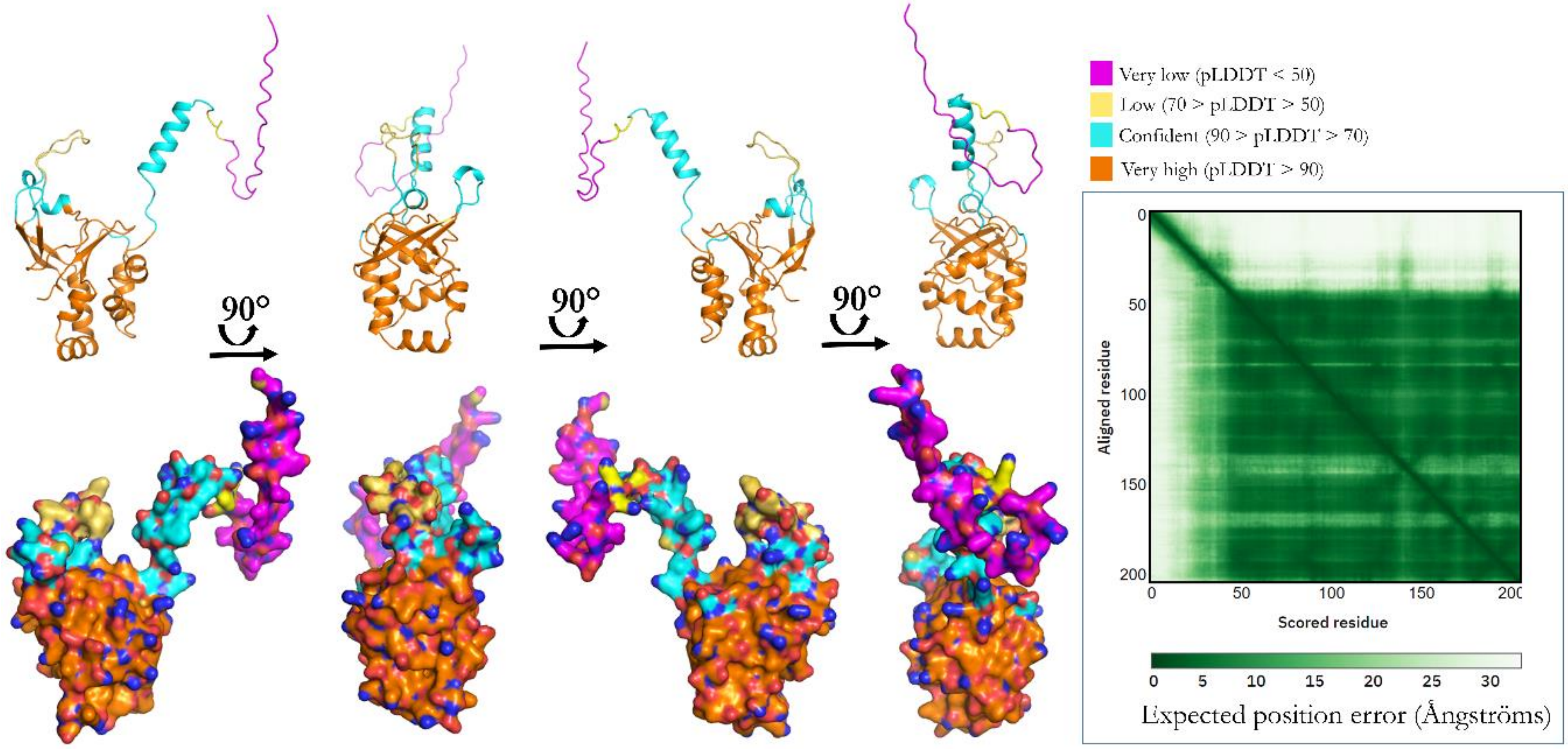

Figure S7. The ab-initio predicted structure of HDPpf (UniProt: Q8IL04) is depicted in this figure. The structure was predicted using DeepMind's AlphaFold algorithm and is represented by a color-coded scheme based on per-residue confidence scores (pLDDT). The structure is shown in both cartoon and surface representations from different orientations. The inset provides information on the predicted alignment error, where the color at each position (x, y) indicates AlphaFold's expected position error at residue x when aligning the predicted and true structures on residue y.

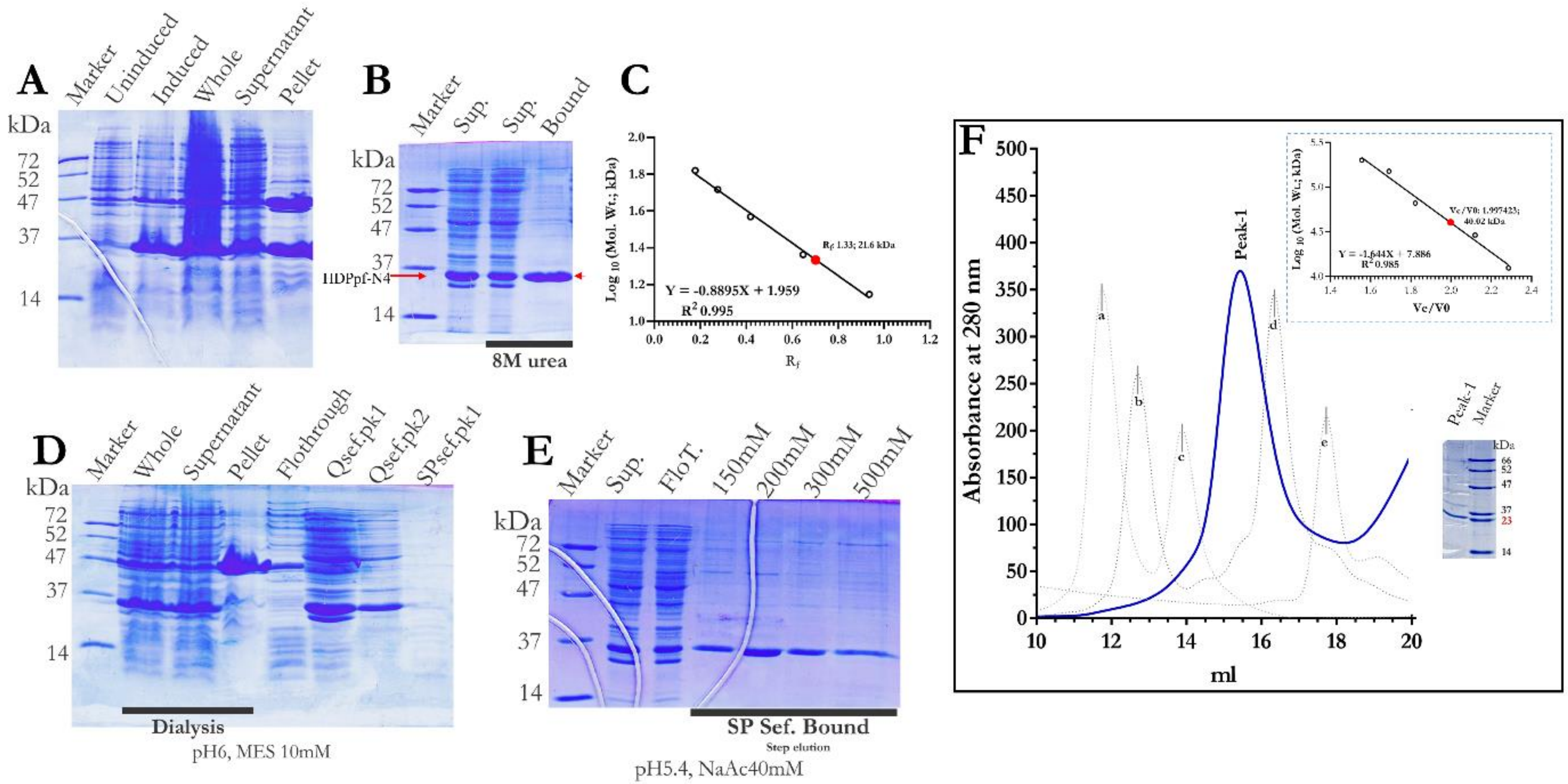

Figure S8. Expression and purification of HDPpf-N4: Coomassie Blue-stained SDS-PAGE (15% W/V) gel of various samples collected during different stages of HDPpf protein purification. **A,** Unindicted, sample exponentially growing culture before IPTG induction; Induced, sample of mature culture after IPTG induction; whole, sample of lysed and homogenized slurry; supernatant, sample of supernatant obtained after centrifugation; pellet, sample of pellet obtained after centrifugation. **B,** Ni-IDA chromatography of HDPpf-N4 protein in presence of 8M urea, **C,** Molecular mass deduction of prominent band in bound lane of image C (red arrow), from Rf value (migration distance of the protein/migration distance of the dye front). The red dot corresponds to the deduced molecular mass of HDPpf-N4. **D,** whole, solubilized ammonium sulphate pellet; supernatant, pellet, sample of supernatant obtained after centrifugation dialyzed ammonium sulphate pellet; flow-through, sample of supernatant unbound Q-SP sepharose beads; Qsef.pk1 & Qsef.pk2, sample bound to Q sepharose beads, SPsef.pk1, sample bound to SP sepharose beads. **E,** Sup., Pooled and dialyzed, bound peaks of Q-sef. Chromatography; FloT., sample of supernatant unbound SP sepharose beads; 150mM, 200mM, 300mM, 500mM, bound peaks eluted by step elution in SP sepharose chromatography. **F,** Gel-filtration elution profile of HDPpf-N4 is represented in blue line. The inset shows the calibration of the Superdex 200 size exclusion column with red dot corresponding to the elution volume of HDPpf-N4. (gel filteration molecular weight markers, a. blue dextran, 2000 kDa, b. alcohol dehydrogenase, 150 kDa., c. albumin bovine serum, 66 kDa, d. carbonic anhydrase, 29 kDa. e. Cytochrome c, horse heart,12.4kDa).

### *Suplimentary information*

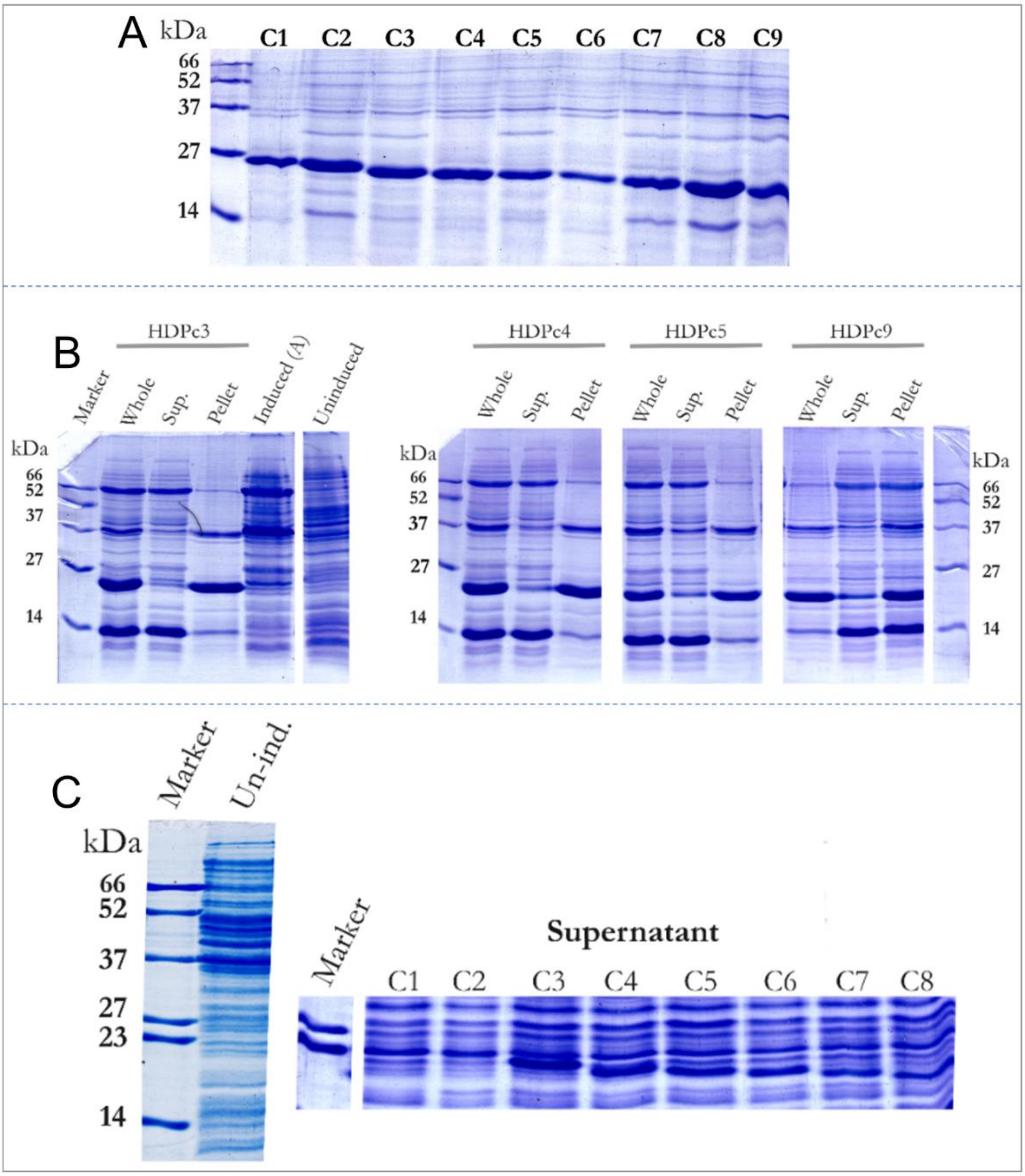

Figure S9. **A,** Coomassie Blue-stained SDS-PAGE (15% W/V) gel of pellet fractions of various constructs of HDPpf with N terminal truncations and C-terminal His tags. **B,** Coomassie Blue-stained SDS-PAGE (15% W/V) gel of various fractions collected during co-expression of HDP C3, C4, C5 and C9 constructs along with pKJE7. (Sup.; Supernatant, induced (A); fraction collected after induction with L-arabinose co-inducer only). **C,** Coomassie Blue-stained SDS-PAGE (15% W/V) gel of various fractions collected during co-expression of HDP constructs (C1-9) along with pGro7. (Un-ind.; un-induced fraction)

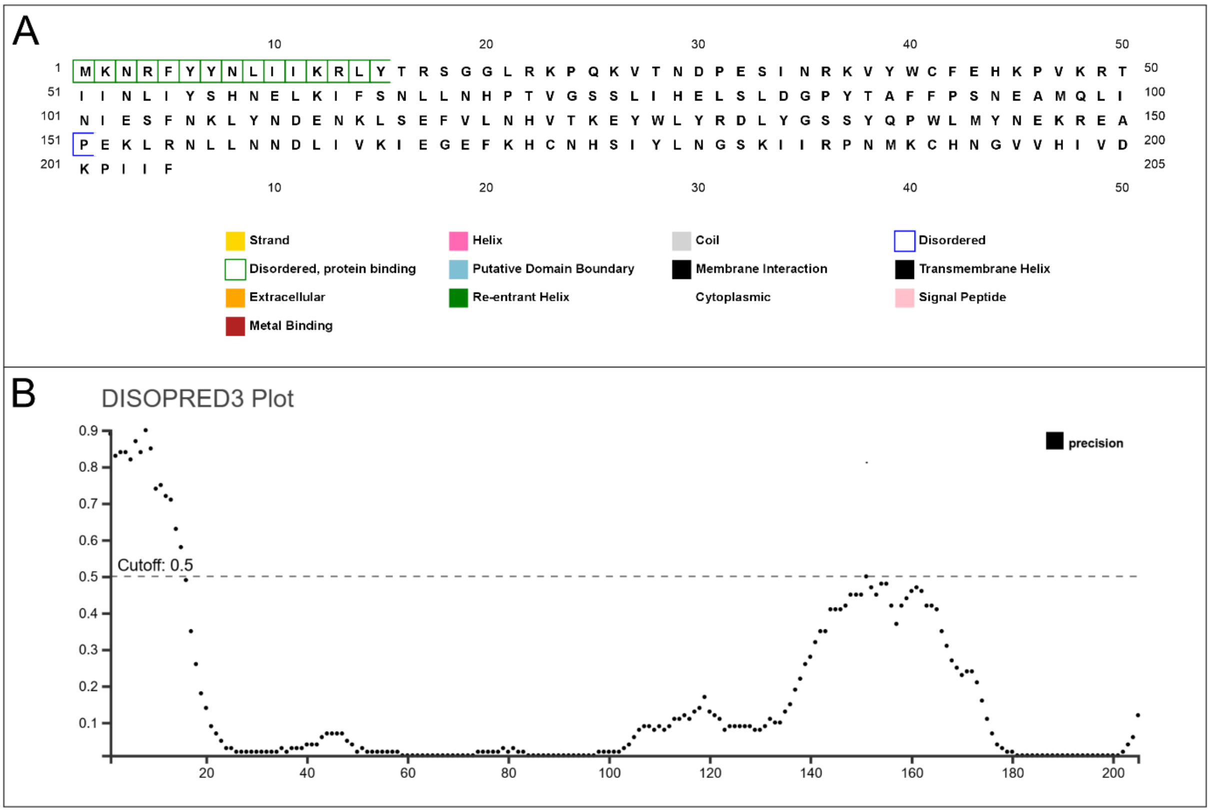

Figure S10. *Prediction of intrinsically disordered regions in Plasmodium falciparum Heme Detoxification Protein (HDPpf) using DISOPRED3.* **A,** *Residue-by-residue disorder annotation of HDPpf.* This panel displays the primary sequence of HDPpf, with colour coded annotated predicted by DISOPRED3. **B,** *Disorder probability plot for HDPpf.* This graph presents the disorder probability across the HDPpf sequence. The X-axis indicates residue number, while the Y-axis shows the probability of disorder (0.0 to 1.0). A red horizontal line at 0.5 marks the classification cutoff—residues above this line are predicted to be disordered.
